# Pathological Mechanisms of Motor Dysfunction in Familial Danish Dementia: Insights from a Knock-In Rat Model

**DOI:** 10.1101/2025.04.15.649002

**Authors:** Arnab Choudhury, Metin Yesiltepe, Tammaryn Lashley, Vishal Singh, Tao Yin, Anllely Fernandez, Luciano D’Adamio, Hyung Jin Ahn

## Abstract

Familial Danish Dementia (FDD) is a rare autosomal dominant neurodegenerative disorder caused by a mutation in the integral membrane protein 2B (*ITM2b*) gene. Clinically, FDD is characterized by cerebral amyloid angiopathy (CAA), cerebellar ataxia, and dementia. Notably, FDD shares several neuropathological features with Alzheimer’s disease (AD), including CAA, neuroinflammation, and neurofibrillary tangles. In this study, we investigate the pathological mechanisms linking CAA, white matter damage, and motor dysfunction using a recently developed FDD knock-in (FDD-KI) rat model. This model harbors the Danish mutation in the endogenous rat *Itm2b* gene, along with an *App* gene encoding humanized amyloid-β (Aβ). Our analysis revealed substantial vascular Danish amyloid (ADan) deposition in the cerebellar subpial and leptomeningeal vessels of FDD-KI rats, showing an age-related increase comparable to that observed in human FDD patients. Additionally, vascular Aβ deposits (Aβ-CAA) were present in FDD-KI rats, but Aβ-CAA patterns showed some differences between species. Motor function assessments in FDD-KI rats demonstrated age-accelerated motor deficits and gait abnormalities, mirroring the clinical characteristics of FDD patients. To further explore the mechanisms underlying these deficits, we examined cerebellar pathology and found age-related myelin disruption and axonal fiber loss, consistent with postmortem human FDD pathology. Cerebellar demyelination appeared to be driven by neuroinflammation, marked by increased microglial/macrophage activation in response to vascular amyloid deposition. Additionally, we observed extravascular fibrinogen leakage, indicating widespread vascular permeability in both white and gray matter, with fibrinogen deposits surrounding amyloid-positive vessels in aged FDD-KI rats and postmortem FDD cerebellum. These findings suggest that this FDD-KI rat model is the first animal model to recapitulate key neuropathological features of human FDD patients, including both ADan- and Aβ-type CAA, neuroinflammation, and white matter lesions—pathologies that may underlie the motor and gait impairments seen in the disease.

## Introduction

Cerebral amyloid angiopathy (CAA) is a prevalent cerebral vessel disease characterized by the accumulation of amyloid peptides around cerebral blood vessels [2, 7, 25]. CAA occurs in both hereditary (familial) and sporadic forms and is present in approximately 80% of Alzheimer’s disease (AD) patients. It is associated with various cerebrovascular events, including cerebral infarction, intracerebral hemorrhages (ICH), microbleeds, and white matter (WM) damage, and contributes to age-related cognitive decline [12, 49, 67]. Despite its prevalence, the precise pathogenic mechanism of CAA remains incompletely understood. In addition to cognitive decline, CAA has been associated with motor dysfunction, as many familial and sporadic CAA patients experience motor deficits and abnormal gait [6, 48]. These motor impairments are closely linked to white matter damage in both cortical and subcortical regions [51, 52]. Furthermore, recent findings suggest that CAA pathology may contribute to cerebellar atrophy, potentially leading to gait disturbances [24]. However, despite this clinical evidence, the underlying mechanisms connecting CAA, white matter damage, and motor dysfunction remain poorly understood, necessitating further investigation.

While CAA is primarily associated with β-amyloid (Aβ), the major amyloid peptide found in Alzheimer’s disease (AD) patients, several other forms of CAA have been identified, including CAA with Danish amyloid (ADan) and CAA with British amyloid (ABri). ADan and ABri amyloids arising from *ITM2B* mutations that cause Familial Danish Dementia (FDD) and Familial British Dementia (FBD), respectively [71, 72]. These variants share the common feature of amyloid deposition in cerebral blood vessels but differ in their genetic origins and biochemical properties. Understanding these distinct forms of CAA can help elucidating the broader mechanisms underlying CAA-associated cerebrovascular pathologies, identifying shared pathogenic pathways, and developing therapeutic strategies with the potential to broadly target and mitigate CAA.

*ITM2B* encodes BRI2, a type II membrane protein that is synthesized as an immature precursor. This precursor undergoes cleavage at the C-terminus by proprotein convertases, generating the mature BRI2 protein and a soluble 23-amino acid C-terminal fragment, Bri23 [11]. In FDD, a decamer duplication insertion before the stop codon in *ITM2B* results in a frameshift mutation, producing a mutant BRI2 protein with an additional 11 amino acids at the C-terminus. Instead of Bri23, proprotein convertase cleavage of this mutant BRI2 generates the 34-amino acid amyloidogenic peptide known as ADan [72]. ADan forms amyloid fibril deposits in the brain parenchyma as well as around blood vessels, resulting in CAA [23, 66]. Notably, CAA with Aβ (Aβ-CAA) is also present in the brains of FDD patients, emphasizing the complex interplay of amyloidogenic proteins in disease pathology. Clinically, FDD is characterized by cataracts, progressive cerebellar ataxia, and dementia [72]. Neuropathologically, FDD shares several features with AD, including widespread amyloid angiopathy in parenchymal and leptomeningeal vessels, ischemic stroke, white matter hyperintensities (WMH), and tau pathology [23, 72].

*ITM2b* has important functions in the CNS. Studies on *Itm2b-KO* and conditional *Itm2b-KO* mice, have shown that BRI2 has a cell autonomous physiological function in synaptic transmission and plasticity in glutamatergic neurons at both presynaptic and postsynaptic termini[73]. BRI2 has a dual anti-amyloidogenic function, reducing both Aβ production and Aβ aggregation. It has been found that BRI2 binds to APP in cis, thereby reducing APP cleavage and Aβ production[35–37, 60]. Additionally, the extracellular domain of BRI2 includes a BRICHOS domain that inhibits or delays Aβ aggregation[10, 61]. These activities of BRI2 on APP processing and Aβ aggregation are supported by evidence that APP and APP processing play a role in long-term synaptic plasticity and memory deficits in FDD knock-in (KI) and FBD-KI mice [57–60]. More recent studies have shown that *ITM2B* expression is highest in microglia [1, 75], which are strongly linked to AD. These findings suggest that BRI2 plays a key role in dementia pathogenesis by regulating APP, as well as through its direct effects on neuronal and microglial function.

Although studies in FDD-KI mice have provided valuable insights, this model has limitations, as it does not fully replicate the neuropathological features observed in human FDD patients [20]. In contrast, the transgenic mouse model for FDD—generated by randomly inserting multiple copies of a human ITM2B mini-gene carrying the Danish mutation under neuron-specific promoters—developed ADan deposits and neuroinflammation, but showed no Aβ deposition or motor symptoms (40, 41). The discrepancy between KI and transgenic models may be due to differences in ADan production, with transgenic models expected to generate much higher ADan levels than FDD-KI mice. Additionally, in mice, the presence of endogenous murine Aβ, which is less prone to amyloid aggregation compared to human Aβ, may hinder the formation of ADan pathology.

To address this potential issue, we studied an FDD-KI rat model carrying the Danish mutation (*Itm2b^D/D^*) along with humanized *App* alleles (*App^h/h^*), introduced via genome editing [62, 63, 74]. This model expresses a mutant *Itm2b* gene under the control of its natural regulatory elements, preventing the excessive mutant protein production seen in transgenic models. Additionally, the *App^h^* allele enables the rat *App* gene to produce APP physiologically while generating human Aβ instead of rat Aβ. This design ensures a more physiologically relevant system for studying disease pathology and progression. Building on insights from previous FDD studies, we hypothesize that the FDD-KI rat serves as a valuable model for studying the relationship between CAA, white matter damage, and motor impairment. To explore this, we assessed age-dependent CAA pathology, focusing on vascular ADan and Aβ deposition in the cerebellum and compared these findings with human FDD postmortem brain tissue. Furthermore, we examined the impact of vascular amyloid deposition on white matter integrity and neuroinflammation, which may contribute to motor dysfunction. This study provides insights into the mechanisms linking cerebellar vascular amyloid deposition, white matter lesions, and motor deficits in AD and related dementias.

## Materials and methods

### Animals

We used FDD-KI (*Itm2b*^D/D^: *App*^h/h^) rat model which has an FDD-associated frameshift mutation (*Itm2b*^D/D^) and carries the *App* gene with the humanized Aβ sequence (*App*^h/h^)[74]. The FDD mutation consists of ten nucleotides duplication one codon before the normal stop codon and a frameshift in the *Itm2b* sequence generating a precursor protein 11 amino acids larger than normal wildtype protein. The rats were housed in an under controlled environment with a 12-hour dark/light cycle, a temperature of 22 ± 2°C, and had free access to food and water. After breeding, the pups were genotyped for FDD mutation by polymerase chain reaction with the following specific forward (F): FAATGTGGAAATTATGGGGTGGAT and reverse (R) primers: RGATGAAGAGACAGTGAAGCCCTG followed by Sanger sequence analysis. Littermate controls (*Itm2b*^W/W^: *App*^h/h^) which carries a wild-type *Itm2b* genes (*Itm2b*^W/W^) and the *App* gene with the humanized Aβ sequence (*App*^h/h^) were used in the entire study. All the experiments were performed according to policies on the care and use of laboratory animals in the Ethical Guidelines for Treatment of Laboratory Animals of the National Institute of Health. All the procedures were described and approved by the Rutgers Institutional Animal Care and Use Committee (IACUC).

### Tissue processing and immunofluorescence

Rats were transcardially perfused with 0.9% saline plus 2.5% heparin solution. Brains were harvested and embedded in optical cutting temperature (OCT) compound and snap-frozen in cryomold on dry-ice and stored at -80°C. Following freezing, brain tissues were cut with 20 µm thickness of sagittal sections using cryostat (CM1950, Leica Microsystems). Tissue sections were mounted serially on coated glass slide, dried and stored at -80°C for further analysis.

Sections were fixed in freshly prepared chilled methanol: acetone (1:1) or 4% paraformaldehyde (PFA) for 20 min at room temperature. For ADan and Aβ staining, sections were fixed in 4% PFA, washed in phosphate buffer saline (PBS) and incubated with 95% formic acid (Sigma Aldrich) for 2 min at room temperature. For delipidation, sections were washed in PBS and submerged in 100% ethanol for 10 min, as required. Sections were then washed in PBS and blocked in PBST (1X PBS + 0.3% Triton X-100) with 5% normal donkey serum for 1 hour at room temperature in humidified condition. Thereafter, sections were incubated with primary antibody diluted in PBST solution (1X PBS + 0.1% Triton X-100) with 5% normal donkey serum at 4°C overnight in humidified chamber. The flowing primary antibody were used: anti-ADan (Ab-1700 polyclonal; 1:500), Anti-amyloid-β, 1-16 (6E10) conjugated to Alexa Fluor 488 (1:250) (BioLegend, 803013), anti-fibrinogen (1:500) (Dako, A008002-2), 3) anti-collagen IV (1:400) (Millipore, AB769), anti-Iba-1 (1:200) (Invitrogen, PA5-18039), anti-GFAP (1:200) (Dako/Agilent Technology, Z0334), anti-MMR/CD206 (R&D Systems, AF2535). Sections were washed three times in PBST (PBS + 0.05% TritonX-100) for 5 minutes with gentle shaking condition and incubated with secondary antibody, diluted in PBST (1X PBS + 0.1% Triton X-100) with 5 % donkey serum for 1 hour at room temperature in humidified condition. The secondary antibody used were donkey anti-rabbit Alexa Fluor 488, donkey anti-goat Alexa Fluor 405, donkey anti-goat Alexa Fluor 488 (1:1000). After secondary antibody incubation, sections were washed 3 times with PBST (1X PBS+0.05% Triton X-100) for 5 minutes on shaker. Tissue sections were incubated with 0.3% Sudan black B (Sigma, 199664) mixed 70% ethanol for 1 minute to minimize lipofuscin autofluorescence and quickly washed with 70% ethanol for 20 seconds, followed by PBS washing, 3 times for 5 minutes on shaker. Sections were then mounted with Vectashield^®^ antifade mounting medium (Vector Labs, H-1000-10) and covered with cover slip.

### Myelin and axonal staining

To assess the myelin integrity in cerebellum white matter, Luxol Fast blue (LFB) staining (Abcam) was performed according to manufacturer’s instruction. Briefly, fresh frozen rat sagittal sections were fixed in 4% PFA and incubated with LFB solution for 24 hours at room temperature. Sections were then submerged in lithium carbonate solution to differentiate and then in ethanol to further differentiate. Then sections were washed and dehydrated with absolute alcohol and dried for several hour before mounted on coverslips using VectaMount™ mounting medium (Vector Laboratories, USA; H-5000). For myelin immunofluorescence, sections were fixed in 4% PFA for 20 min at room temperature. Then sections were washed in PBS and submerged in 100% ethanol for 10 min for delipidation. Sections were washed in PBS three times for 5 min each and permeabilized with 2% normal donkey serum diluted in 1% Triton X-100 for 30 min at room temperature. After washing with PBS, sections were blocked with Super Block blocking buffer diluted in PBS (Thermo Fisher Scientific) for another 30 min at room temperature. Tissues sections were incubated with Myelin Basic Protein (MBP) (Cell Signaling Technology, 78896; 1:200) and Neurofilament heavy chain (1:1500) (Millipore, AB5539) diluted in 5% normal donkey serum with 1X PBS + 0.1% Triton X-100 at 4°C for overnight. After washing with PBST, sections were incubated in secondary antibodies anti-rabbit Alexa Fluor 488 and donkey anti-chicken Alexa Fluor 555 respectively for 1 hour at room temperature. Sections were washed in PBST and treated with 0.3% Sudan Black B solution 70% ethanol for 1 minute to minimize lipofuscin autofluorescence. Following PBS washing sections were mounted with Vectashield^®^ antifade mounting medium (Vector Labs, H-1000-10).

### Image analysis and quantification

The stained sagittal rat brain sections were analyzed and processed using Nikon Ti2 Eclipse fluorescence microscope equipped with a motorized stage, a digital CMOS camera (DS-Qi2, Nikon) and Lumencor Spectra light engine. To minimize biased stereology, fluorescence images of whole sagittal brain sections were acquired using tile scanning (14 x 22 field of view) with an 15% overlap, utilizing Nikon CFI Plan Apochromat Lambda 20x/0.75 NA objective. Each set of stained sagittal sections was processed under identical gain, SOLA power, and LUT settings using NIS element AR software (Nikon). The whole sagittal sections were analyzed for ADan neuropathology. The following workflow was used in Fiji image J (NIH) to analyze the stained images. The total area of ADan and Aβ deposits were analyzed by measuring stained area as percentage of total cerebellum area. The number of ADan and Aβ positive CAA-laden vessels were manually counted in each cerebellum area. The number of extravascular fibrinogen deposits were determined by manually counted fibrin(ogen) deposits that were extravasated at least 50% outside vessel area. Similarly, the total area of GFAP or Iba-1-positive activated microglia was analyzed by measuring the astrocyte and mihcroglia-stained area as a percentage of total area of cerebellum. The LFB, MBP and neurofilament heavy chain (NFH) stained area was analyzed using area threshold and particle analysis to determine the stained area as a percent of total cerebellum area. At least 3 sections per each rat were used in all image analysis and the researcher was blinded to the genotype of each rat.

### Behavioral assessment for motor coordination and gait

#### Rotarod test

The accelerating rotarod (IITC Life Sciences Inc. USA) test is a classical sensory-motor task to assess motor coordination and balance skills of animals by measuring the ability of the animal to stay and run on the accelerated rod [53]. Rats were habituated for four days before performing test. Rats were subjected to three trials per day for 3 consecutive days allowing them to train for the test the interval between trials in a day was 30 min. Each 600 sec trial began with a 30-s acclimation period at 4 rpm followed by gradual acceleration to a maximum of 30 rpm over the remaining phase of the trial. Performance was measured as the latency to fall for rat from the rod and the average was calculated.

#### Catwalk test

The quantitative gait analysis related to cerebellar dysfunction was performed using Catwalk XT system (Noldus Information Technology, Wageningen, The Netherlands). The system consists of a 1.3m black corridor situated perpendicular to a glass plate with green LED light illuminated inside, forming a walkway that animals freely traverse one side to other. During the walk, intensely illuminated animal paw prints were captured by a high-speed video camera positioned underneath the walkway. The experiments were performed at 7- and 13-month aged rats according to the manufacturer’s instructions and focused on gait parameters previously identified as ataxic gait [8, 30, 42, 65]. Rats were habituated to the experimental room for four days and allowed to cross the walkway for 2-3 min voluntarily to the Catwalk instrument for 3-4 days to become familiar with the apparatus. On the day of test run, rats were tested by freely crossing the walkway for three compliant runs, described as walking across the corridor without stopping, turn around and change direction. Minimum and maximum run duration was set as 0.5 and 5 sec, respectively with 60% maximum speed variation. For all the rats tested, the detection setting was used as (Camera Gain: 20.20 dB, Green Intensity Threshold: male-0.13; Female-0.10, Red Ceiling Light: 17.8, and Green Walkway Light: 16.0). For analysis, gait parameters were automatically generated using Catwalk XT software (Version 10.6) after each paw print being manually classified and labeled as LF (Left Front), LH (Left Hind), RF (Right Front), and RH (Right Hind) paws. Gait parameters were categorized as run characteristic and kinetic parameter, temporal parameters, spatial parameters and interlimb coordination parameters. All the behavioral experiments and analysis were performed by experimenter blinded to the genotype of animals.

### Postmortem human brain tissues

Postmortem human tissue samples were obtained from Queen Square Brain Bank, Queen Square Institute of Neurology, University College London, United Kingdom. We obtained fresh frozen and formalin fixed cerebellum tissues of FDD cases. We included non-demented (ND) control cerebellum tissues for immunohistochemical analysis. For all the brain tissue, case demographics were summarized in Supplementary Table 1.

### Fresh frozen human tissue processing and immunofluorescence

Fresh frozen postmortem human cerebellum tissues were cut with 20 μm thick sections using cryostat (CM1950, Leica Microsystems). Tissue sections were mounted on coated glass slide, dried and stored at -80°C for further analysis. For immunofluorescence staining,sections were fixed in 4% paraformaldehyde (PFA) for 20 min at room temperature or in freshly prepared chilled methanol: acetone (1:1) for 5 min at -20°C. For ADan and Aβ staining, sections were fixed in 4% PFA, washed in phosphate buffer saline (PBS) and incubated with 95% formic acid (Sigma Aldrich) for 2 min at room temperature. Sections were then washed in PBS and blocked in PBST (1X PBS + 0.3% Triton X-100) with 2% normal donkey:2% normal horse serum (1:1) for 1 hour at room temperature in humidified condition. Thereafter, sections were incubated with primary antibody diluted in PBST solution (1X PBS + 0.1% Triton X-100) with 2% normal donkey:2% normal horse serum (1:1) at 4°C overnight in humidified chamber. The flowing primary antibody were used: anti-ADan (Ab-1700 polyclonal; 1:500), Anti-amyloid-β, 1-16 (6E10) conjugated to Alexa Fluor 488 (1:250) (BioLegend, 803013), anti-fibrinogen (1:500) (Dako, A008002-2), 3) anti-collagen IV (1:400) (Millipore, AB769). Sections were washed three times in PBST (PBS+0.05% TritonX-100) for 5 minutes with gentle shaking condition and incubated with secondary antibody, diluted in PBST (1X PBS + 0.1% Triton X-100) with 2 % normal donkey: 2: normal horse serum (1:1) for 1 hour at room temperature in humidified condition. The secondary antibody used were donkey anti-rabbit Alexa Fluor 488, donkey anti-goat Alexa Fluor 405 (1:1000). After secondary antibody incubation, sections were washed 3 times with PBST (1X PBS+0.05% Triton X-100) for 5 minutes on shaker. Tissue sections were incubated with 0.3% Sudan black B (Sigma, 199664) mixed 70% ethanol for 1 minute to minimize lipofuscin autofluorescence and quickly washed with 70% ethanol for 20 seconds, followed by PBS washing, 3 times for 5 minutes on shaker. Sections were then mounted with Vectashield^®^ antifade mounting medium (Vector Labs, H-1000-10) with cover slip and images were acquired using Nikon Ti2 eclipse fluorescence microscope with NIS elements AR software.

### Formalin fixed tissue histology and immunohistochemistry

For neurofilament cocktail and CR3-43 staining in FDD patient cerebellum, paraffin embedded sections were deparaffinized in xylene and rehydrated in varying concentrations of alcohol. The endogenous peroxidase activity was blocked with 0.3% H_2_O_2_ in methanol for 10 mins. Sections were pressure cooked for 10 mins in citrate buffer pH 6.0, then incubated in 10% non-fat milk to block any non-specific staining. Sections were incubated in the primary antibodies against HLA class II histocompatibility antigen (clone CR3-43, Dako 1:150) and neurofilament cocktail (monoclonal 1:20, ICN Pharmaceuticals) for 1 hour at room temp. This was followed by several washes in PBS and then treatment with either biotinylated anti-mouse (Dako 1:200) or biotinylated anti-rabbit (1:200) for 30 mins, ABC (Dako) for 30 mins. The peroxidase activity was developed with diaminobenzidine/ H_2_O_2_ solution, counterstained in Meyers Hematoxylin or combined with Congo Red. The stained slides were digitized using an Evident Slide scanner (VS100).

### Statistical analysis

All numerical values were presented in individual point graphs are mean ± SEM and plotted using GraphPad Prism 10 (GraphPad Software, San Diego, CA). In all the experiments, statistical significance was determined using Student’s *t* test or two-way ANOVA and post hoc pairwise t-test with the Bonferroni correction. Comparison of each day training curves in rotarod test was analyzed using two-way ANOVA with repeated measure and Bonferroni post hoc test.

## Results

### ADan deposits within cerebellar vasculature of FDD-KI rat

We assessed ADan-CAA in the brain tissue of FDD-KI (*Itm2b*^D/D^: *App*^h/h^) rats using immunohistochemistry. Immunofluorescence analysis with ADan antibody on whole sagittal rat brain sections revealed that ADan deposition was exclusively localized to the cerebellum in FDD-KI rats, with no detectable deposits in other brain regions (Fig. 1a and Supplementary Fig. 1). Given this cerebellum-specific distribution, we focused all subsequent neuropathological assessments on this region. Further analysis revealed that ADan deposits were primarily localized within subpial and leptomeningeal vessels, with a lesser presence in capillaries. Additionally, non-vascular plaques were predominantly found in the molecular layer of the cerebellum in *Itm2b*^D/D^: *App*^h/h^ rats compared to *Itm2b*^W/W^: *App*^h/h^littermate rats (Fig. 1a, b). No ADan deposition were detected in the cerebellum of *Itm2b*^D/W^: *App*^h/h^ rat. Notably, the extent of ADan deposition occurs in the FDD-KI rats increased with age. At 7-month of age, *Itm2b*^D/D^: *App*^h/h^rats exhibited a trend toward increased ADan deposition in the cerebellum. However, by 13 and 16 months, the cerebellar ADan-positive area was significantly greater in *Itm2b*^D/D^: *App*^h/h^ rats compared to *Itm2b*^W/W^: *App*^h/h^ rats (Fig. 1c, d). To further assess the amyloidogenic nature of ADan deposits, we performed co-staining with Congo Red (CR). Interestingly, most CR-stained areas were co-localized with ADan deposits in the cerebellum of *Itm2b*^D/D^: *App*^h/h^ rat (Supplementary Fig. 1). To validate these neuropathological findings in the FDD-KI model, we also conducted ADan immunofluorescence analysis on postmortem cerebellar tissues from FDD patients and control subjects. ADan depositions were highly abundant within subpial large vessels and medium parenchymal vessels and with lesser extent in capillaries in the molecular layer of cerebellar cortex area in FDD patients compared to ND controls (Fig. 1e, f). ADan positive parenchymal plaques were also present within the cerebellar cortex (Fig. 1f). The distribution of ADan deposits around blood vessels and within the parenchyma in the *Itm2b*^D/D^: *App*^h/h^ rat cerebella closely mirrored the pathology observed in FDD patient cerebella (Fig. 1b, f). These findings confirm that the ADan pathology in the FDD-KI rat model closely recapitulates the neuropathological features observed in FDD patients.

**Fig. 1.**
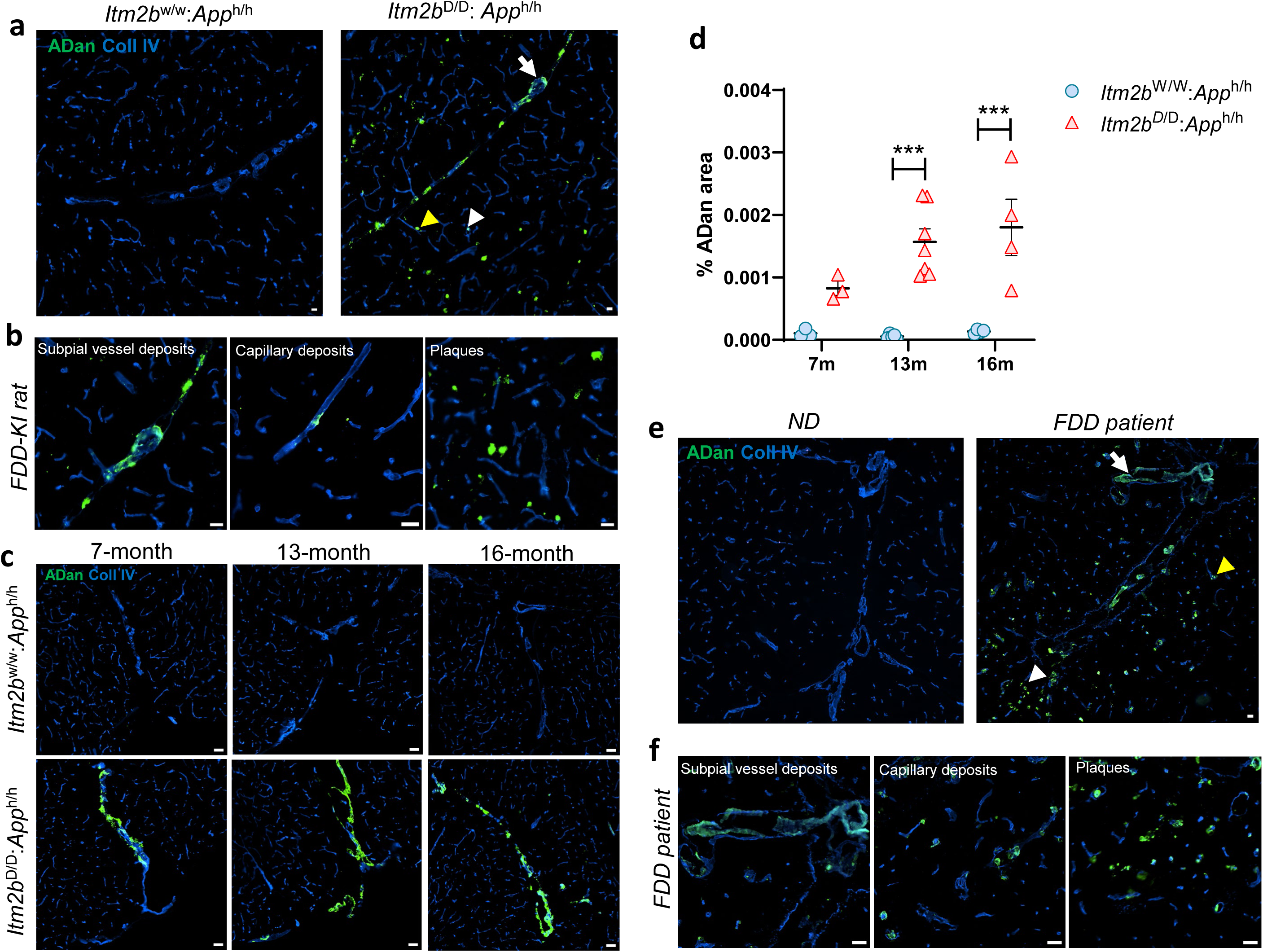
ADan pathology in FDD-KI rat and patient carrying *Itm2b* mutation in the cerebellum. (a) ADan deposits in subpial and leptomeningeal vessels (white arrow), capillaries (yellow arrowhead), and parenchymal plaques (white arrowhead) in the cortical area of cerebellum of *Itm2b*^D/D^: *App*^h/h^ rats compared to *Itm2b*^W/W^: *App*^h/h^ rats. (b) Magnified images showing ADan-laden leptomeningeal vessels, capillaries and plaques in *Itm2b*^D/D^: *App*^h/h^ rat cerebella. (c) Immunofluorescence images showing age-related ADan deposition in the cerebellum of *Itm2b*^D/D^: *App*^h/h^ and *Itm2b*^W/W^: *App*^h/h^ rats. (g) ADan-deposited areas showed an age-dependent increase in the cerebellum of *Itm2b*^D/D^: *App*^h/h^ rats at 7, 13, and 16 months compared to *Itm2b*^W/W^: *App*^h/h^ rats. (n = 3-7). (e) ADan deposition was observed in subpial vessels (white arrow), capillaries (yellow arrowhead), and independent plaques (white arrowhead), predominantly in the molecular layer of the cerebellar cortex in FDD patients. No ADan immunoreactivity found in ND control cerebellum. (f) Magnified images showing ADan deposition in leptomeningeal vessels, capillaries and ADan-positive independent plaques in FDD patient cerebella. Data were analyzed by two-way ANOVA with the Bonferroni post hoc test and shown as mean ± SEM; ***P < 0.001. Scale bars = 50 μm, Coll IV= Collagen type IV.

### Vascular A**β** deposition in FDD cerebellum

To assess the Aβ-CAA pathology in the cerebellum of FDD-KI rat model, immunostaining was performed using Aβ (6E10) antibody along with blood vessel marker collagen IV (Coll IV). Independent Aβ immunolabeling revealed that Aβ (6E10) deposits were primarily observed in parenchymal medium sized vessel and capillaries in the cerebellum white matter *Itm2b*^D/D^: *App*^h/h^ rats compared to *Itm2b*^W/W^: *App*^h/h^ rats. An age-dependent increase in Aβ-CAA vessels was observed in the deep white matter of 7-, 13- and 16-month-old *Itm2b*^D/D^: *App*^h/h^rats compared to *Itm2b*^W/W^: *App*^h/h^ rats (Fig. 2a, b). No subpial or leptomeningeal Aβ-CAA or Aβ plaques were detected in the cerebellum of *Itm2b*^D/D^: *App*^h/h^ rat. In FDD patient cerebellum, Aβ-CAA was observed in fewer medium-sized vessels and capillaries in the deep white matter (Fig. 2c, right panel), while it was predominantly found in subpial large vessels, with no deposition ND control cerebellum (Fig. 2c, left panel). These results suggest that while Aβ predominantly deposits in subpial large vessels in the cerebellum of FDD patients, it primarily accumulates in medium-sized vessels and capillaries in the white matter of FDD-KI rat cerebellum.

**Fig. 2.**
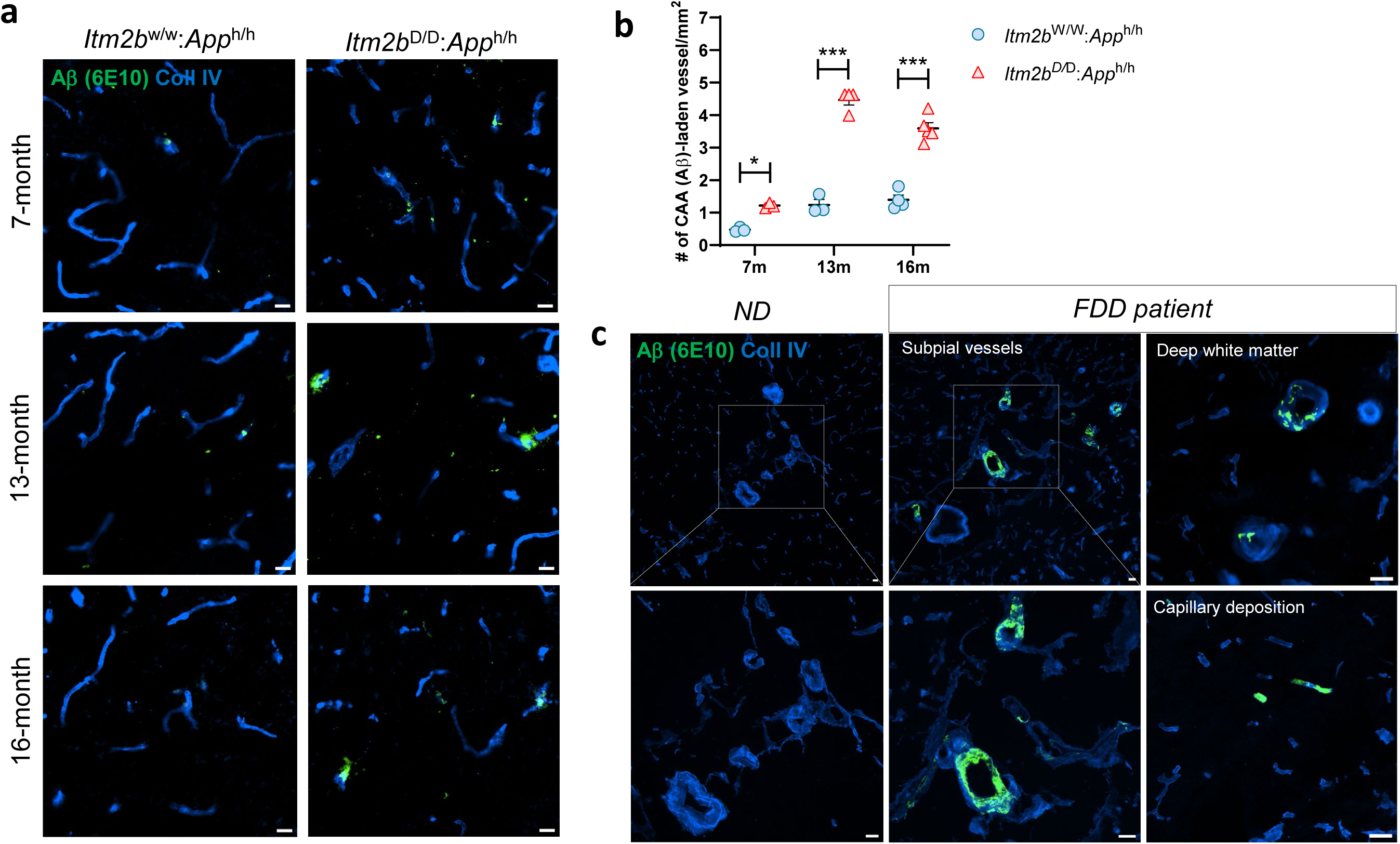
Aβ immunofluorescence in FDD-KI rat and patient cerebellum. (a) Age-dependent Aβ (6E10) deposition in the capillaries and medium-sized vessels within the white matter of cerebellum of *Itm2b*^D/D^: *App*^h/h^ rats compared to *Itm2b*^W/W^: *App*^h/h^ rats. (b) Aβ-positive CAA-laden vessels increase in the cerebellum of *Itm2b*^D/D^: *App*^h/h^ rats at 7, 13, and 16 months compared to *Itm2b*^W/W^: *App*^h/h^ rats. (n = 3-6). (c) Aβ (6E10) deposition observed in the cerebellar subpial large vessels of FDD patient compared to control subjects. With lesser extent Aβ deposited in medium-sized vessels and capillaries within the deep white matter of FDD patients (right panels). Data were analyzed by two-way ANOVA with the Bonferroni post hoc test and shown as mean ± SEM; * P< 0.05, ***P < 0.001. Scale bars = 50 μm, Coll IV= Collagen type IV.

### Motor impairment and abnormal gait behavior in FDD-KI rats

Clinical studies indicated that FDD patients develop cerebellar ataxia, characterized by a loss of coordination and balance while walking, typically in their fourth decade of life [23]. Given that cerebellar vascular lesions often lead to motor deficits [24, 27], we assessed cerebellar-influenced motor behaviors in FDD-KI rats to evaluate the potential association between vascular amyloid deposition and age-related motor impairment. We employed the Rotarod test, a widely used method for assessing somatosensory coordination and motor learning in rodents [18]. At 7-month age, the motor learning performance of *Itm2b*^D/D^: *App*^h/h^ rats on the accelerating rotarod was similar to that of wild-type rats over a 3-day trial (Fig. 3a). However, at 13 months of age, significant differences in the latency to fall were observed on day 3 of the trial between *Itm2b*^D/D^: *App*^h/h^ rats and *Itm2b*^W/W^: *App*^h/h^ rats at 13-month age (Fig. 3b), indicating a deficit in balance and motor learning in *Itm2b*^D/D^: *App*^h/h^ rat. We further evaluated gait parameters of FDD-KI rats using the Catwalk XT automated gait analysis system at 7 and 13 months of age. At 7 months, no significant differences were observed in any gait parameters between *Itm2b*^D/D^: *App*^h/h^ and *Itm2b*^W/W^: *App*^h/h^ rats (Fig. 3c). However, at 13 months age, *Itm2b* ^/^ : *App* ^/^ rats exhibited a significant decrease in right paw stance duration, measured as the time a front or hind paw remained on the glass without support from the contralateral paws (Fig. 3d). We also analyzed the percentage of single-paw and three-paw support during walking, which reflects interlimb coordination patterns—specifically, the configuration of paws on the ground during stance, often altered in ataxic gait [26]. At 13-months, *Itm2b*^D/D^: *App*^h/h^ rats also showed a significant increase in single-paw support, accompanied by a compensatory decrease in three-paw support, compared to *Itm2b*^W/W^: *App*^h/h^ rats (Fig. 3e, f). Taken together, these findings suggest that FDD-KI rats develop age-dependent cerebellar-related motor coordination deficits and ataxic gait abnormalities, mirroring the phenotype observed in FDD patients.

**Fig. 3.**
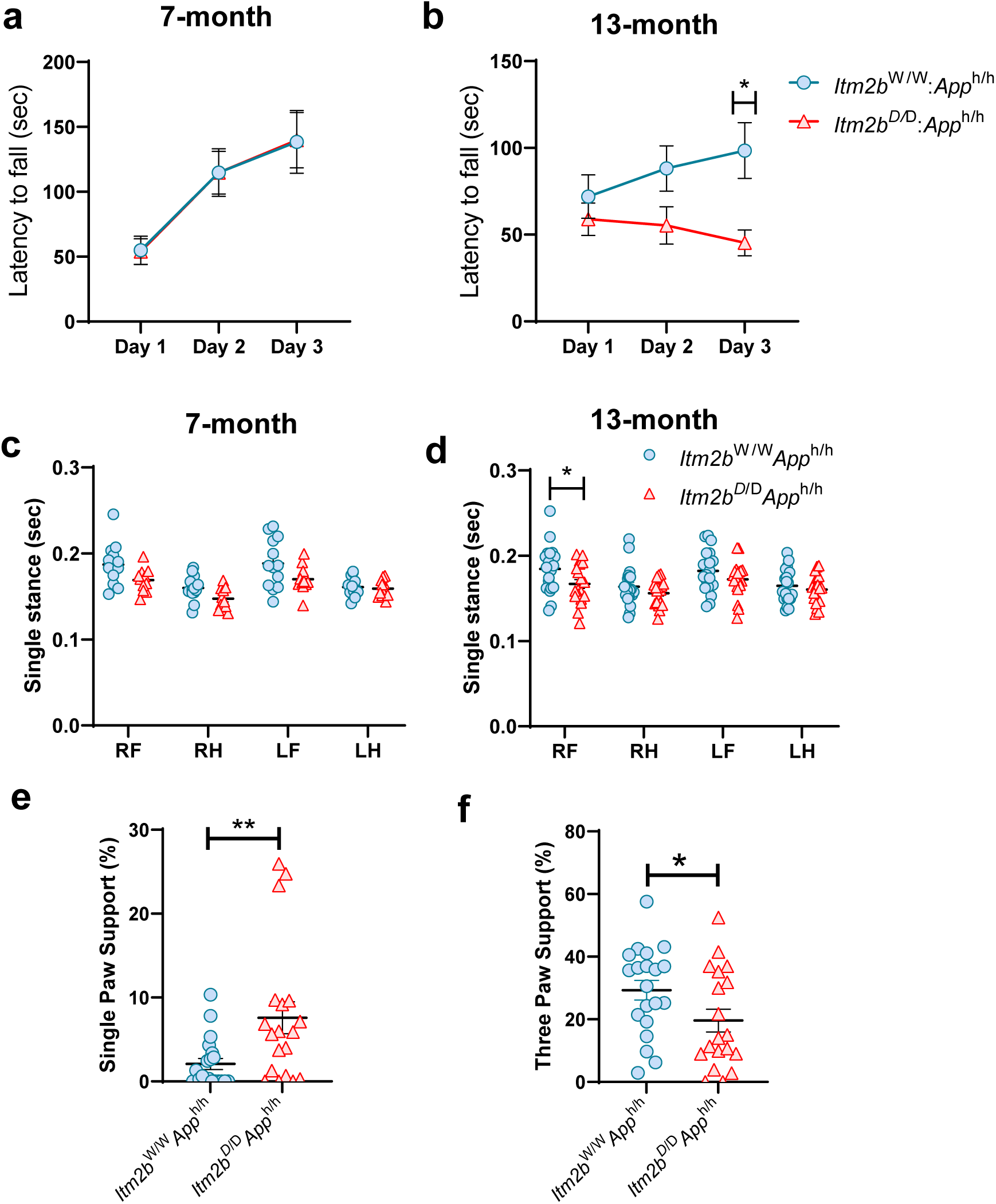
Altered motor coordination and abnormal gait in FDD-KI rat. (a, b) Motor learning and coordination were measured using accelerating rotarod in 7- and 13-month-old *Itm2b*^D/D^: *App*^h/h^ rats and compared to *Itm2b*^W/W^: *App*^h/h^ rats. The latency of fall from the rotating rod was analyzed for three trials per day for three consecutive days (n = 9-10). (c, d) Gait parameters were measured by the Catwalk XT gait analysis system. Graphs showing paw specific single stance in 7- and 13-month-old *Itm2b*^D/D^: *App*^h/h^ rats compared to *Itm2b*^W/W^: *App*^h/h^ rats. Single stance represents percent time the front or hind paw is standing on the walkway without the contralateral paw. (e, f) Graphs showing percentage of single and three paw supports during walking in 7- and 13-month-old *Itm2b*^D/D^: *App*^h/h^ rats compared to *Itm2b*^W/W^: *App*^h/h^ rats (n = 19-20). Data were analyzed by Student’s *t* test or two-way ANOVA with the Bonferroni post hoc test and shown as mean ± SEM; * P< 0.05, *P<0.01. RF = Right Front, LF = Left Front, RH = Right Hind, LH = Left Hind.

### Cerebellar demyelination and axonal damage in FDD-KI rat

White matter lesions and lacunar infarcts associated with cerebral small vessel disease are believed to impair connections between key motor regions, leading to motor and gait disturbances [17, 45, 67]. In both familial and sporadic CAA, vascular amyloid deposition, followed by cerebral microbleeds and hemorrhage, is associated with increased white matter lesions in cortical and subcortical areas [19, 54, 55]. In FDD patients, MRI imaging showed white matter changes, and neuropathological evaluation confirmed myelin damage with axonal loss in the deep cerebellar white matter [23]. To investigate whether motor deficits and cerebral amyloid deposition in the cerebellum is associated with white matter demyelination in FDD-KI rats, we analyzed myelin basic protein (MBP), a marker for myelination used to determine structural integrity of axonal tract, in the cerebellum of *Itm2b*^W/W^: *App*^h/h^ and *Itm2b*^D/D^: *App*^h/h^ rat. At 7-month age, no significant differences in MBP area were found in the cerebellum white matter of *Itm2b*^D/D^: *App*^h/h^ compared to *Itm2b*^W/W^: *App*^h/h^ rat (Fig. 4a, c). However, MBP expression was significantly reduced both in the cerebellar foliar white matter and sparse myelin fibers in the granule cell layer of 13-month-old *Itm2b*^D/D^: *App*^h/h^ rat than *Itm2b*^W/W^: *App*^h/h^ rat (Fig. 4b, d). Furthermore, Luxol Fast Blue (LFB) staining in 13-month also revealed a significant reduction in myelin density in the cerebellum white matter of *Itm2b*^D/D^: *App*^h/h^ rat (Supplementary Fig. 2a, b). To determine whether myelin loss leads to axonal damage, we assessed neurofilament heavy chain (NFH) immunostaining in the cerebellum. As expected, NFH-stained axonal tracts integrity showed no changes in 7-month-old *Itm2b*^D/D^: *App*^h/h^ rat (Fig. 4e, g). However, at 13 months, *Itm2b*^D/D^: *App*^h/h^ rats exhibited a significant reduction in NFH immunoreactivity in both the cerebellar white and gray matter compared to *Itm2b*^W/W^: *App*^h/h^ rat (Fig. 4f-h).

**Fig. 4.**
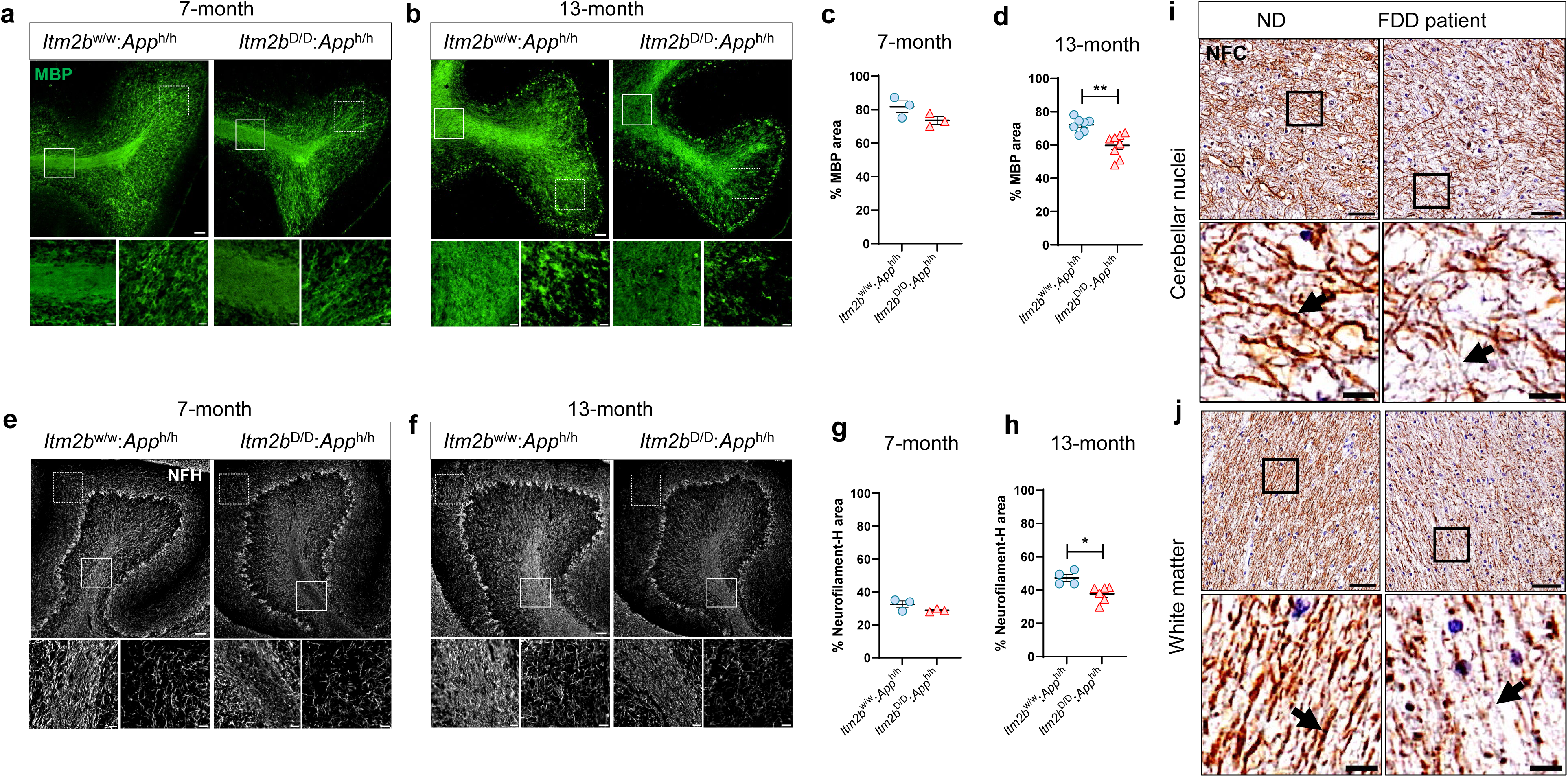
Cerebellar demyelination and axonal loss in aged FDD-KI rat. (a, b) Representative immunofluorescence images of myelin basic protein (MBP) showing demyelination in the cerebellum of *Itm2b*^D/D^: *App*^h/h^ and *Itm2b*^W/W^: *App*^h/h^ rats at 7- and 13-month age. Enlarged views of the regions marked by solid boxes (foliar white matter) and dotted boxes (areas with sparse fibers) are shown in the lower panels. (c, d) MBP stained area was quantified as percentage of total cerebellum area at 7- and 13-month age. (n = 3-8) (e, f) Representative images showing neurofilament heavy chain (NFH) immunoreactive axonal fibers in the cerebellum of *Itm2b*^D/D^: *App*^h/h^ and *Itm2b*^W/W^: *App*^h/h^ rats at 7- and 13-month age. Enlarged views of the regions marked by solid boxes (foliar white matter) and dotted boxes (axon fibers in the molecular layer) are shown in the lower panels. (g, h) Quantification of NFH stained area as percentage of total cerebellum area in 7- and 13-month age. (n = 3-8) (I, j) Representative immunohistochemistry images showing neurofilament cocktail (NFC) stained axonal fibers (black arrows) in the foliar white matter and cerebellar nuclei region of postmortem FDD patient and ND control subjects. Enlarged views of the regions marked by solid boxes are shown in the lower panels. Data were analyzed by Student’s *t* test and shown as mean ± SEM; * P< 0.05, **P< 0.01. Scale bars = 50 μm; 10μm.

We next investigated whether the pathological characteristics of axonal damage observed in the FDD-KI rat model were also present in an FDD patient. Neurofilament cocktail staining of the cerebellum from FDD patients revealed significant axonal fiber damage in both the white matter and cerebellar nuclei compared to ND controls (Fig. 4i, j). Axonal tracts in ND controls appeared continuous, whereas those in FDD patients were fragmented or discontinuous (indicated by black arrows in Fig. 4i and 4j), suggesting significant axonal damage in the cerebellum. Similarly, in our FDD-KI rat model, NFH immunolabeling showed comparable features of axonal damage, with thinner or disintegrated tracts observed in both the white matter and cerebellar nuclei of 13-month-old *Itm2b*^D/D^: *App*^h/h^ rats (Supplementary Fig. 2c). In contrast, *Itm2b*^W/W^: *App*^h/h^ rat exhibited a higher density of intact axonal tracts (Supplementary Fig. 2c). Overall, these findings suggest that the FDD-KI rat model exhibits age-related myelin damage and axonal loss in the cerebellum, closely mirroring the pathological features observed in postmortem cerebellar tissue from FDD patients.

### Microglial activation in white matter demyelination of FDD-KI rats

Pro-inflammatory activation of microglia and macrophages is a key driver of demyelination [13, 34]. In this study, we aimed to investigate the mechanisms underlying white matter demyelination in the cerebellum of the FDD-KI rat model and explore whether neuroinflammation in response to vascular amyloid deposition contributes to this process. Iba-1 immunofluorescence at 7-month age revealed slight but not significant increase in Iba-1 positive cells in cerebellum of *Itm2b*^D/D^: *App*^h/h^ rat compared to wild-type littermates rats (Fig.5a, b). At 13-month age, significant increase in Iba-1 positive cells were found in cerebellum covering predominantly white matter and cerebellar nuclei in *Itm2b*^D/D^: *App*^h/h^ rat (Fig. 5a, b). GFAP positive astrocyte reactivity was not significantly increased in both 7- and 13-month age in *Itm2b*^D/D^: *App*^h/h^ rats (Supplementary Fig.3a, b). We further analyzed the perivascular macrophage population in the cerebellum, given their proximity to CAA and crucial role in amyloid clearance [22]. At 13-month age, a significant increase in immunofluorescence area of mannose receptor CD206 was found in cerebellum of *Itm2b*^D/D^: *App*^h/h^ rat compared to wild-type littermates rats (Fig. 5c, d). These CD206-positive perivascular macrophages were predominantly associated with amyloid deposition around subpial and leptomeningeal vessels. To examine microgliosis in FDD patients, we performed immunohistochemistry on postmortem cerebellar tissue using HLA class II histocompatibility antigen (clone CR3/43) to identify activated microglial populations. Combined with Congo Red staining, this analysis revealed that activated microglia were primarily engaged with amyloid deposits within blood vessels or plaques in the cerebellar cortex of FDD patients (Fig. 5e). In contrast, control cerebellar tissue showed no presence of activated microglia in these regions. In the white matter, an increased number of activated microglial cells were observed surrounding Congo Red-positive vessels as well as independently of vessels, compared to ND control tissues (Fig. 5e). These findings suggest that activated microglia may be involved in white matter demyelination, while perivascular macrophages primarily engage with vascular amyloid deposits.

**Fig. 5.**
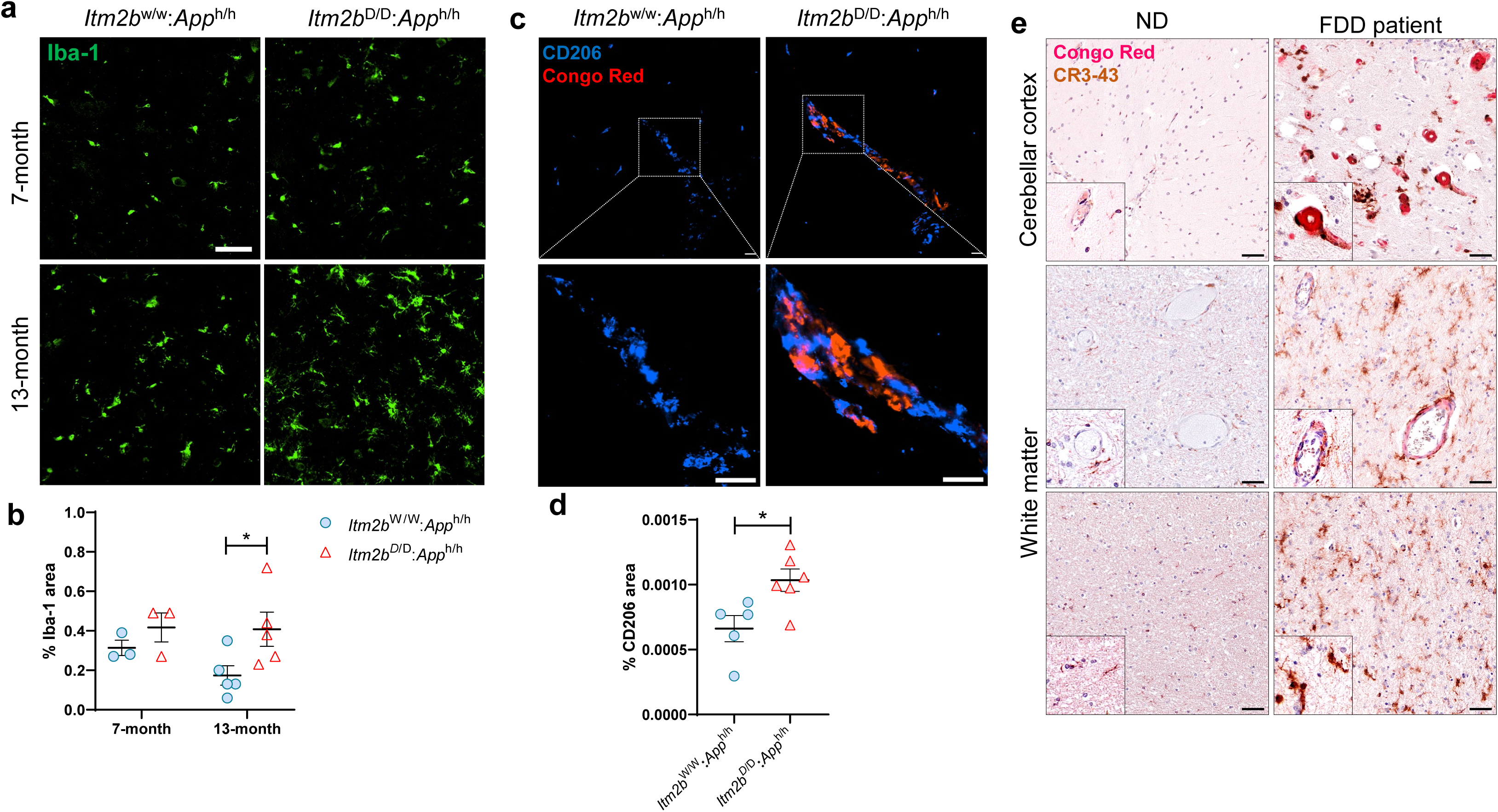
Microglia/macrophage activation and antigen presentation in the FDD cerebellum. (a) Immunofluorescence images of Iba-1 showing microglia/macrophage activation in the cerebellum of *Itm2b*^D/D^: *App*^h/h^ and *Itm2b*^W/W^: *App*^h/h^ rats at 7- and 13-month age. Scale bars = 50 μm (b) Quantification of Iba-1stained area shown as percentage of total cerebellum area at 7- and 13-month age. (n = 3-5). (c) Representative images showing perivascular macrophage specific marker CD206 along with Congo Red staining in cerebellum of *Itm2b*^D/D^: *App*^h/h^ and *Itm2b*^W/W^: *App*^h/h^ rats at 13-month age. Scale bars = 50 μm; 10μm (d) Quantification of CD206 area shown as percentage of total cerebellum area at 13-month age. (n = 5-6). (e) Immunohistochemistry images of HLA class II histocompatibility antigen (CR3-43) positive reactive microglial/macrophage counterstained with Congo Red in postmortem FDD brain compared with ND control tissue. CR3-43 positive microglia showing association with Congo Red stained blood vessels in cerebellar cortex. In the white matter, CR3-43 reactive microglia were either vessel associated, or non-vessels associated. Scale bars = 50 μm. Data were analyzed by Student’s *t* test or two-way ANOVA with the Bonferroni post hoc test and shown as mean ± SEM; * P< 0.05.

### Increase of extravasated fibrinogen in cerebellum of FDD patient and FDD-KI rat

Elevated levels of perivascular and parenchymal fibrinogen deposition have been observed in the brains of AD and Hereditary Cerebral Amyloid Angiopathy (HCAA) patients, as well as in rodent models of AD [5, 9]. This contributes to blood-brain barrier (BBB) permeability, neuroinflammation, and motor dysfunction [5, 9]. To investigate whether neuroinflammation in the cerebellum is associated with increased BBB permeability in response to CAA, we analyzed fibrinogen leakage from blood vessel using immunofluorescence in FDD-KI rat and wild-type cerebellum tissue. At 7-monoth age, number of extravasated fibrinogen spots were not significantly changed while at 13-month age, we observed a significant increase in extravasated fibrinogen spots throughout white and grey matter of cerebellum of *Itm2b*^D/D^: *App*^h/h^ rats compared to *Itm2b*^W/W^: *App*^h/h^ rats (Fig. 6a-c).

**Fig. 6.**
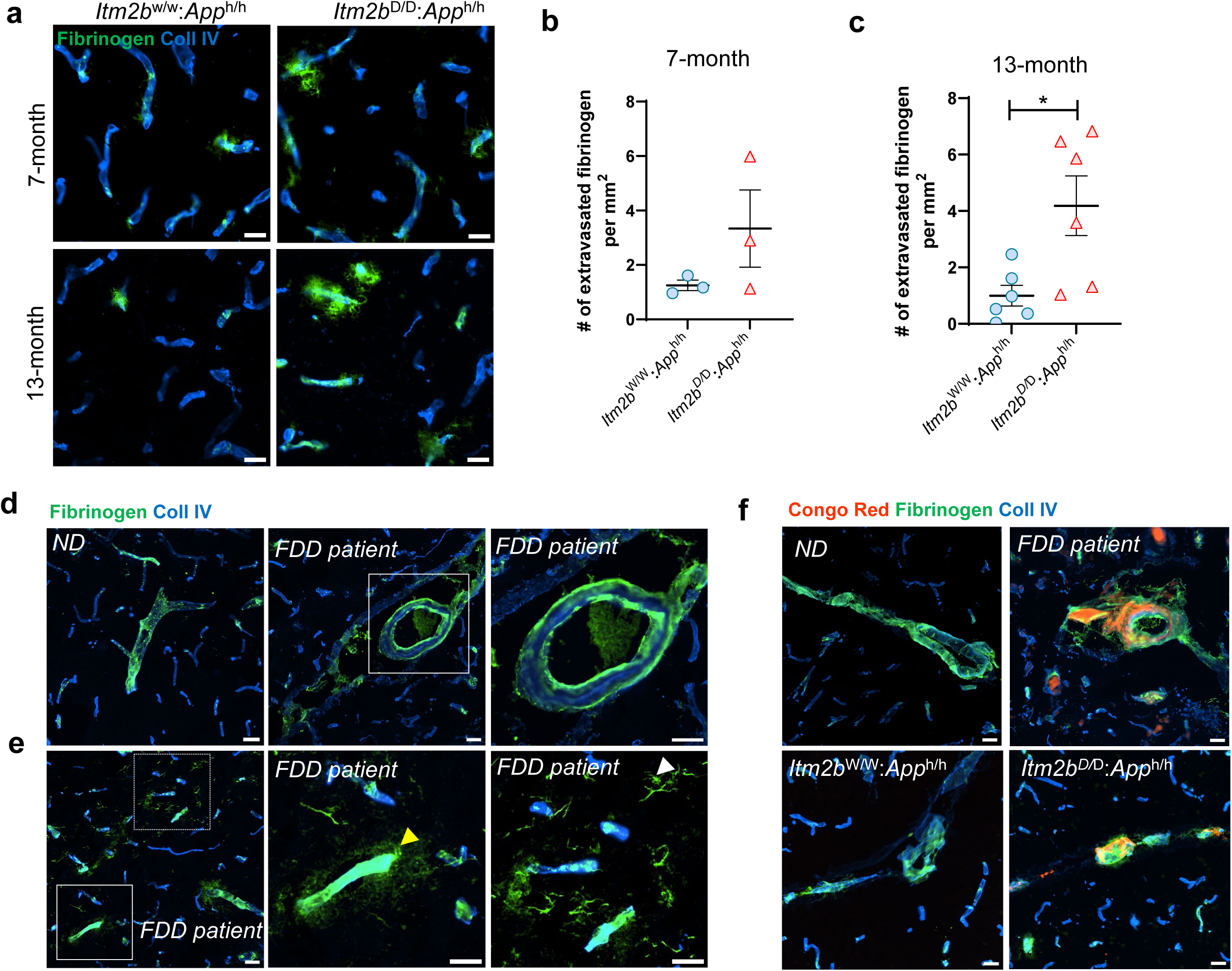
Extravasated fibrinogen in FDD patient and knock-in rat cerebellum: (a) Fibrinogen extravasation in cerebellum of *Itm2b*^D/D^: *App*^h/h^ rats compared to *Itm2b*^W/W^: *App*^h/h^ rats at 7- and 13-month age. (b, c) Number of extravasated fibrinogen was measured in the whole cerebellum at 7- and 13-months (n = 3-6) (d) Fibrinogen immunoreactivity observed in the outer layer and extravascular space of subpial vessels in the cerebellum of FDD patient. High magnification images of medium-sized vessels and capillaries shows extravascular fibrinogen leakage into the surrounding parenchyma (yellow arrowhead). Fibrinogen-positive glial-like cell morphology in the white matter area (white arrowheads). In the ND control subjects, fibrinogen is majorly inside the vessels. (e) Extravasated fibrinogen in the Congo Red positive cerebellar leptomeningeal vessels in FDD patient and *Itm2b*^D/D^: *App*^h/h^ rats compared to ND controls and *Itm2b*^W/W^: *App*^h/h^ rats respectively. Data were analyzed by Student’s *t* test and shown as mean ± SEM; * P< 0.05; Coll IV= Collagen type IV; Scale bars = 50 μm.

In FDD patient cerebellum, distinct features of fibrinogen extravasation were observed, including leakage from subpial large vessels, showing fibrinogen immunoreactivity in the outer layer and the extravascular space, indicating extravascular fibrinogen deposition compared to control cerebellum (Fig. 6d). In line with previous studies reporting parenchymal and white matter fibrinogen deposition in Alzheimer’s disease (AD) and multiple sclerosis (MS) [21, 31, 39, 68], we also observed fibrinogen leakage into the surrounding parenchyma in small arteries and capillaries, along with fibrinogen-positive glial-like cell morphology in the white matter of the cerebellum in FDD patients (Fig. 6e). Furthermore, we examined whether fibrinogen extravasation occurs within ADan-deposited vessels. Immunolabeling with Congo Red (CR) in postmortem cerebellum revealed that fibrinogen extravasation occurred predominantly within the CR-positive leptomeningeal vessels, with a similar co-deposition of ADan and fibrinogen observed in the 13-month-old *Itm2b*^W/W^: *App*^h/h^ rat cerebellum (Fig. 6f). As expected, fibrinogen was primarily confined to the intravascular space in the cerebellum of ND controls and *Itm2b*^W/W^: *App*^h/h^ cerebellums (Fig. 6f). Taken together, these results suggest that vascular amyloid deposition and microglial activation may be linked to the increased extravasation of fibrinogen in the FDD cerebellum.

## Discussion

Several neurodegenerative diseases, including AD and FDD, are characterized by abnormal vascular and parenchymal amyloid protein accumulations, white matter lesions, and neuroinflammation. These shared features likely result from common underlying mechanisms, which may contribute to abnormal behavior symptoms, including motor dysfunction. In this study, we demonstrated that the presence of familial Danish mutation with humanized Aβ in FDD-KI rats led to prominent age-dependent increase in ADan deposition in cerebellar subpial and leptomeningeal vessels, closely mirroring the cerebellar pathology observed in FDD patients. Additionally, an age-related increase in Aβ deposits was detected primarily in the capillaries of cerebellar white matter. Our findings suggest that demyelination and axonal damage in aged FDD-KI rats may be driven by microglial and macrophage activation in response to extravasated fibrinogen. This vascular pathology, combined with myelin loss, likely contributes to impaired motor coordination and ataxia-related gait abnormalities in FDD. Overall, our results highlight a complex interplay between demyelination, axonal loss, and vascular pathology—including CAA and fibrinogen extravasation—that may underlie motor impairments in FDD.

Transgenic mice carrying the Danish mutation (Tg-FDD) have been developed, exhibiting age-dependent increase of ADan-CAA throughout the brain, along with parenchymal diffuse plaques [70]. In Tg-FDD mice, randomly inserted transgene drives the expression of the mutant form of the human *Itm2b* gene under the control of the mouse prion protein (moPrnp) promoter. ADan-CAA is predominantly observed in cerebellar pial vessels, as well as in medium-sized parenchymal and penetrating vessels of the cerebrum. Another Tg-FDD mouse model, in which the ADan precursor protein was expressed under the control of a cosmid-based Syrian hamster prion protein expression cassette, showed age-related vascular ADan lesions throughout the brain, including the hippocampus, amygdala, and thalamus, but to a lesser extent in the cerebellum [12]. While these transgenic models successfully replicate ADan-associated pathology through overexpression of the mutant *Itm2b* gene, they fail to capture other critical pathological and behavioral features observed in human disease, such as Aβ-CAA, neuroinflammation and cerebellar ataxia. Moreover, because these models rely on artificial promoters, they drive the expression of multiple copies of the mutant gene in a spatiotemporal pattern that differs from the endogenous allele, limiting their translational relevance.

To overcome these limitations, a knock-in (KI) approach was developed to more accurately model the genetic basis of human FDD. The FDD-KI mouse model, generated by introducing the pathogenic decamer duplication in exon 6 of the *Itm2b* gene, exhibits synaptic plasticity and memory deficits but lacks overt brain pathology or motor impairments [38, 57, 59, 60]. Since FDD-KI mice do not exhibit detectable vascular or parenchymal ADan and Aβ deposition [20], the FDD-KI rat model was developed using CRISPR/Cas-mediated genome engineering to express the mutant *Itm2b* gene under endogenous regulatory control. Given that BRI2 interacts with and modulates APP processing, and human Aβ species exhibit greater pathological potential than their rodent counterparts, a humanized Aβ sequence (*App^h^*) was introduced into FDD-KI rats [62, 63]. This modification enabled the FDD-KI rat model to more effectively replicate the cerebral amyloidosis observed in FDD patients. Indeed, we observed both ADan-CAA and Aβ-CAA in FDD-KI rats, closely resembling amyloid pathology in postmortem cerebellar tissue from FDD patients. However, amyloid pathology appeared more pronounced in human cases than in the rat model, likely due to endpoint postmortem analysis in patients [23, 72]. Notably, FDD-KI rats exhibit not only synaptic deficits [74], but also vascular amyloid pathology, neuroinflammation, and motor dysfunction, whereas FDD-KI mice display synaptic and memory deficits without overt brain pathology or motor impairments. This discrepancy suggests that while the loss of BRI2 function due to the FDD mutation may primarily drive synaptic and memory deficits, the presence of vascular amyloid pathology—particularly in the cerebellum—could be a key contributor to neuroinflammation and motor dysfunction. However, further investigation is needed to validate this hypothesis.

In FDD patients, Aβ deposition was found to colocalize with the ADan peptide in various brain regions, either a similar distribution pattern to ADan or occurring independently. However, Aβ-positive vessels were significantly fewer than ADan-positive ones [23, 66]. To date, no Tg-FDD mouse models have exhibited Aβ-CAA or parenchymal Aβ deposition [12, 70]. In contrast, the FDD-KI model showed Aβ-CAA primarily in capillaries and small vessels within the cerebellar white matter at 7 and 13 months of age. CAA is classified into two forms: type-1, marked by amyloid deposition in capillaries and microvessels, and type-2, characterized by deposits in larger vessels, including leptomeningeal and intracortical small arteries and arterioles [3, 49, 64]. Recently developed transgenic rat models have demonstrated these distinctions, with the rTg-DI model (carrying both the Dutch E693Q and Iowa D694N familial CAA mutations) developing type-1 CAA, while the rTg-D model (harboring only the Dutch Aβ mutation) exhibited type-2 CAA in larger vessels [15, 16]. Interestingly, the FDD-KI rat model developed ADan-CAA, which may be classified as type-2 CAA due to its predominant deposition in subpial and leptomeningeal vessels. Meanwhile, Aβ-CAA observed in the white matter could be categorized as type-1 CAA.

In the early phase of CAA formation, altered BBB permeability increases fibrinogen leakage through endothelial cell layer, resulting in fibrin(ogen) deposition within the perivascular space [28, 33]. In the cerebellar white matter of FDD patient, fibrinogen-immunoreactive glial-like cells were present, likely due to phagocytosis of fibrinogen by glial cells, as reported in previous studies [28, 39]. Earlier reports suggest that fibrinogen leakage induces perivascular clustering of microglia by interact with them through CD11b-CD18 receptor, resulting in neuroinflammation and contributing to spine elimination and axonal loss [14, 40]. Transgenic CAA rat model with progressive type-1 CAA or capillary CAA demonstrated strong perivascular neuroinflammation, whereas in type-2 CAA with more fibrillar amyloid deposited within large vessels showed less or no perivascular glial activation [16, 76]. In FDD-KI rat cerebellum, increased microglial activation was observed in the white and grey matter where Aβ-CAA and fibrinogen extravasation was more prominent. Coincided with previous reports, less Iba-1-positive microglia or macrophage were observed near areas of ADan-CAA deposition in pial and subpial vessels. Perivascular macrophage populations have been shown to involve in clearance of amyloid and phagocytic activity in type-2 CAA [22]. Immunofluorescence staining with CD206 have indicated that perivascular macrophages were activated and closely associated with vascular amyloid, possibly due to their proximity to deposited ADan in the subpial large vessels in FDD-KI rat cerebellum. In the FDD patient hippocampus and subiculum, CR3/43 positive microglia recognized the MHC-class II antigen, densely populated around CAA-bearing blood vessels [23]. Similarly, our results in postmortem FDD showed CR3/43 positive microglia or macrophage clustered near amyloid deposited blood vessels in grey and white matter of the cerebellum. This neuroinflammatory response in the FDD cerebellum appears to be primarily associated with combined effects of CAA and fibrinogen extravasation.

Extravascular fibrinogen deposition promotes inflammation induced demyelination in the white matter of MS and experimental autoimmune encephalomyelitis (EAE) animal models [31, 44, 50]. Widespread demyelination in the cerebellar cortex has also been documented in MS patients [29]. Microglial activation and pro-inflammatory signaling are essential in driving demyelination, followed by myelin clearance [13, 34]. Our previous study showed demyelination and axonal loss in the striatum following Aβ deposition and fibrinogen extravasation. [5]. The present study showed an age-related increase in demyelination, indicated by reduced MBP immunoreactive myelin structure in the cerebellum of FDD-KI rat. In FDD patients, white matter lesions have been reported using MRI [23]. Notably, reduced NFH-positive axonal density in the cerebellum of FDD-KI rat were comparable to axonal damage observed in FDD patient. These findings suggest that increased BBB permeability may lead to fibrinogen extravasation and microglial activation, collectively creating an inflammatory environment that drives myelin damage and contributes to axonal loss in the FDD cerebellum. Recent evidence suggests a link between white matter damage and motor as well as gait impairments in patients with CAA [47, 51, 52]. Preclinical models of cerebellar white matter lesions and Purkinje cell degeneration demonstrate abnormal motor coordination and gait impairment [26, 41]. Additionally, toxin-induced demyelination in both the corpus callosum and cerebellum results in motor dysfunction in MS mouse model [69]. In this study, the FDD-KI rat model demonstrated age-related deficits in motor coordination and learning, as assessed by the rotarod test. The cerebellum plays a critical role in motor coordination and balance, and impairments in these functions are often associated with cerebellar ataxia. Patients with cerebellar ataxia struggle with executing well-coordinated, target-directed movements, particularly in limb coordination [4, 46]. Cerebellar myelin deficit is associated to motor dysfunction in rotarod test [32, 69]. Consistent with these findings, our rat model exhibited progressive motor coordination impairments on the rotarod test, potentially due to cerebellar white matter damage. Beyond movement coordination, cerebellar ataxia also affects gait and balance, leading to inaccurate foot placement and stride variability [49]. Mouse models of ataxia with cerebellar atrophy show gait impairments, including increased three-paw support and altered cadence [56]. Ataxia mouse model with cerebellar atrophy have showed gait impairment including support on three paw and altered cadence [43]. Similarly, the 13-month-old FDD-KI rats exhibited reduced three-paw support with a compensatory increase in single-paw support, indicating abnormal gait patterns.

## Conclusion

In this study, we present a novel FDD-KI rat model that closely replicates the neuropathology observed in human FDD patients. This model exhibits age-related ADan deposition predominantly in subpial and leptomeningeal vessels of the cerebellum, along with Aβ accumulation primarily in capillaries within the cerebellar white matter. These vascular amyloid deposits may promote fibrinogen extravasation and neuroinflammatory responses, potentially compromising myelin integrity and leading to axonal damage, which likely underlies motor dysfunction and abnormal gait behavior. Further research is needed to elucidate the mechanisms underlying inflammation-induced demyelination in FDD progression. Overall, the FDD-KI rat model provides a valuable platform for studying CAA progression and offers key insights into the potential links between motor dysfunction, CAA, and white matter pathology in FDD.

## Acknowledgements

We thank Dr. Ruben Vidal (Indiana University School of Medicine) for providing ADan antibodies. This work was supported by NIH Grants NS104386 (to H.J.A.), AG078245 (to H.J.A.), and AG033007 (to L. D.). The Queen Square Brain Bank is supported by the Reta Lila Weston Institute of Neurological Studies, UCL Queen Square Institute of Neurology

---

**Supplementary Fig 1 Adan-CAA colocalized with Congo Red staining in FDD-KI rat cerebellum** (a) ADan immunofluorescence and Congo red-stained whole cerebellum of FDD-KI rat. Scale bars = 1000 μm. (b) Magnified images show ADan-positive subpial and leptomeningeal vessels were colocalized with Congo Red-stained vessels in the *Itm2b*^D/D^: *App*^h/h^ rat. Scale bars = 500 μm

**Supplementary Fig 2 Reduction in myelin structure and axonal fibers in FDD-KI rat cerebellum.** (a) Luxol Fast Blue (LFB) stained cerebellum of 13-month-old *Itm2b*^D/D^: *App*^h/h^ rats compared to *Itm2b*^W/W^: *App*^h/h^ rats. Scale bar = 500 μm (b) LFB-stained area was quantified as percentage of total area of the cerebellum at 13-month age (n = 4-5). Data were analyzed by Student’s *t* test and shown as mean ± SEM; * P< 0.05. (c) Representative immunofluorescence images showing characteristics of neurofilament heavy chain (NFH) stained axonal fibers in the cerebellar nuclei and white matter region of *Itm2b*^D/D^: *App*^h/h^ and *Itm2b*^W/W^: *App*^h/h^ rats at 13-month age. Scale bars = 50 μm.

**Supplementary Fig 3 Astrocyte reactivity in the cerebellum of FDD-KI rat**. (a) Immunofluorescence images of GFAP showing reactive astrocytes in the cerebellum of *Itm2b*^D/D^: *App*^h/h^ and *Itm2b*^W/W^: *App*^h/h^ rats at 7- and 13-month age. (b) Quantification of GFAP stained area shown as percentage of total cerebellum area at 7- and 13-month age. (n = 3-6). There is no difference in the area of reactive astrocytes between *Itm2b*^D/D^: *App*^h/h^ and *Itm2b*^W/W^: *App*^h/h^ rats. Data were analyzed by two-way ANOVA with the Bonferroni post hoc test and shown as mean ± SEM; * P< 0.05. Scale bars = 50 μm.

**Supplementary Table 1.** Case demographics of post-mortem human FDD patients and control brains used in immunohistochemical analyses.

| Cases # | AAD | Gender | Clinical Diagnosis | Tissue Type |
| --- | --- | --- | --- | --- |
| 1 | 43 | M | FDD | FFPE |
| 2 | 60 | F | FDD | FFPE |
| 3 | 58 | M | FDD | FFPE |
| 4 | 52 | M | FDD | FFPE/FF |
| 5 | 57 | M | FDD | FF |
| 6 | 96 | F | Control | FFPE/FF |
| 7 | 82 | F | Control | FFPE/FF |
| 8 | 85 | M | Control | FFPE/FF |
| 9 | 77 | F | Control | FFPE/FF |

AAD age at death; FDD Familial Danish Dementia; F female; M male; FFPE formalin-fixed paraffin-embedded; FF fresh frozen.

## References

1 Arber C, Casey JM, Crawford S, Rambarack N, Yaman U, Wiethoff S, Augustin E, Piers TM, Price M, Rostagno A et al (2024) Microglia contribute to the production of the amyloidogenic ABri peptide in familial British dementia. Acta Neuropathol 148: 65 Doi 10.1007/s00401-024-02820-z

2 Arvanitakis Z, Leurgans SE, Wang Z, Wilson RS, Bennett DA, Schneider JA (2011) Cerebral amyloid angiopathy pathology and cognitive domains in older persons. Ann Neurol 69: 320–327 Doi 10.1002/ana.22112

3 Attems J, Jellinger K, Thal DR, Van Nostrand W (2011) Review: sporadic cerebral amyloid angiopathy. Neuropathol Appl Neurobiol 37: 75–93 Doi 10.1111/j.1365-2990.2010.01137.x

4 Bastian AJ, Zackowski KM, Thach WT (2000) Cerebellar ataxia: torque deficiency or torque mismatch between joints? J Neurophysiol 83: 3019–3030 Doi 10.1152/jn.2000.83.5.3019

5 Berk-Rauch HE, Choudhury A, Richards AT, Singh PK, Chen ZL, Norris EH, Strickland S, Ahn HJ (2023) Striatal fibrinogen extravasation and vascular degeneration correlate with motor dysfunction in an aging mouse model of Alzheimer’s disease. Front Aging Neurosci 15: 1064178 Doi 10.3389/fnagi.2023.1064178

6 Biffi A (2022) Main features of hereditary cerebral amyloid angiopathies: A systematic review. Cereb Circ Cogn Behav 3: 100124 Doi 10.1016/j.cccb.2022.100124

7 Brenowitz WD, Nelson PT, Besser LM, Heller KB, Kukull WA (2015) Cerebral amyloid angiopathy and its co-occurrence with Alzheimer’s disease and other cerebrovascular neuropathologic changes. Neurobiol Aging 36: 2702–2708 Doi 10.1016/j.neurobiolaging.2015.06.028

8 Caballero-Garrido E, Pena-Philippides JC, Galochkina Z, Erhardt E, Roitbak T (2017) Characterization of long-term gait deficits in mouse dMCAO, using the CatWalk system. Behav Brain Res 331: 282–296 Doi 10.1016/j.bbr.2017.05.042

9 Cajamarca SA, Norris EH, van der Weerd L, Strickland S, Ahn HJ (2020) Cerebral amyloid angiopathy-linked beta-amyloid mutations promote cerebral fibrin deposits via increased binding affinity for fibrinogen. Proc Natl Acad Sci U S A 117: 14482–14492 Doi 10.1073/pnas.1921327117

10 Chen G, Andrade-Talavera Y, Tambaro S, Leppert A, Nilsson HE, Zhong X, Landreh M, Nilsson P, Hebert H, Biverstål H et al (2020) Augmentation of Bri2 molecular chaperone activity against amyloid-β reduces neurotoxicity in mouse hippocampus in vitro. Commun Biol 3: 32 Doi 10.1038/s42003-020-0757-z

11 Choi SI, Vidal R, Frangione B, Levy E (2004) Axonal transport of British and Danish amyloid peptides via secretory vesicles. FASEB J 18: 373–375 Doi 10.1096/fj.03-0730fje

12 Coomaraswamy J, Kilger E, Wolfing H, Schafer C, Kaeser SA, Wegenast-Braun BM, Hefendehl JK, Wolburg H, Mazzella M, Ghiso J et al (2010) Modeling familial Danish dementia in mice supports the concept of the amyloid hypothesis of Alzheimer’s disease. Proc Natl Acad Sci U S A 107: 7969–7974 Doi 10.1073/pnas.1001056107

13 Cunha MI, Su M, Cantuti-Castelvetri L, Muller SA, Schifferer M, Djannatian M, Alexopoulos I, van der Meer F, Winkler A, van Ham TJ et al (2020) Pro-inflammatory activation following demyelination is required for myelin clearance and oligodendrogenesis. J Exp Med 217: Doi 10.1084/jem.20191390

14 Davalos D, Ryu JK, Merlini M, Baeten KM, Le Moan N, Petersen MA, Deerinck TJ, Smirnoff DS, Bedard C, Hakozaki H et al (2012) Fibrinogen-induced perivascular microglial clustering is required for the development of axonal damage in neuroinflammation. Nat Commun 3: 1227 Doi 10.1038/ncomms2230

15 Davis J, Xu F, Hatfield J, Lee H, Hoos MD, Popescu D, Crooks E, Kim R, Smith SO, Robinson JK et al (2018) A Novel Transgenic Rat Model of Robust Cerebral Microvascular Amyloid with Prominent Vasculopathy. Am J Pathol 188: 2877–2889 Doi 10.1016/j.ajpath.2018.07.030

16 Davis J, Xu F, Zhu X, Van Nostrand WE (2022) rTg-D: A novel transgenic rat model of cerebral amyloid angiopathy Type-2. Cereb Circ Cogn Behav 3: 100133 Doi 10.1016/j.cccb.2022.100133

17 de Laat KF, Tuladhar AM, van Norden AG, Norris DG, Zwiers MP, de Leeuw FE (2011) Loss of white matter integrity is associated with gait disorders in cerebral small vessel disease. Brain 134: 73–83 Doi 10.1093/brain/awq343

18 Deacon RM (2013) Measuring motor coordination in mice. J Vis Exp: e2609 Doi 10.3791/2609

19 Fotiadis P, Reijmer YD, Van Veluw SJ, Martinez-Ramirez S, Karahanoglu FI, Gokcal E, Schwab KM, Alzheimer’s Disease Neuroimaging Initiative study g, Goldstein JN, Rosand J et al (2020) White matter atrophy in cerebral amyloid angiopathy. Neurology 95: e554–e562 Doi 10.1212/WNL.0000000000010017

20 Giliberto L, Matsuda S, Vidal R, D’Adamio L (2009) Generation and initial characterization of FDD knock in mice. PLoS One 4: e7900 Doi 10.1371/journal.pone.0007900

21 Hainsworth AH, Minett T, Andoh J, Forster G, Bhide I, Barrick TR, Elderfield K, Jeevahan J, Markus HS, Bridges LR (2017) Neuropathology of White Matter Lesions, Blood-Brain Barrier Dysfunction, and Dementia. Stroke 48: 2799–2804 Doi 10.1161/STROKEAHA.117.018101

22 Hawkes CA, McLaurin J (2009) Selective targeting of perivascular macrophages for clearance of beta-amyloid in cerebral amyloid angiopathy. Proc Natl Acad Sci U S A 106: 1261–1266 Doi 10.1073/pnas.0805453106

23 Holton JL, Lashley T, Ghiso J, Braendgaard H, Vidal R, Guerin CJ, Gibb G, Hanger DP, Rostagno A, Anderton BH et al (2002) Familial Danish dementia: a novel form of cerebral amyloidosis associated with deposition of both amyloid-Dan and amyloid-beta. J Neuropathol Exp Neurol 61: 254–267 Doi 10.1093/jnen/61.3.254

24 Horn MJ, Gokcal E, Becker AJ, Das AS, Warren AD, Alzheimer Disease Neuroimaging I, Schwab K, Goldstein JN, Biffi A, Rosand J et al (2022) Cerebellar atrophy and its implications on gait in cerebral amyloid angiopathy. J Neurol Neurosurg Psychiatry: Doi 10.1136/jnnp-2021-328553

25 Iadecola C, Gottesman RF (2018) Cerebrovascular Alterations in Alzheimer Disease. Circ Res 123: 406–408 Doi 10.1161/CIRCRESAHA.118.313400

26 Jaarsma D, Birkisdottir MB, van Vossen R, Oomen D, Akhiyat O, Vermeij WP, Koekkoek SKE, De Zeeuw CI, Bosman LWJ (2024) Different Purkinje cell pathologies cause specific patterns of progressive gait ataxia in mice. Neurobiol Dis 192: 106422 Doi 10.1016/j.nbd.2024.106422

27 Konczak J, Pierscianek D, Hirsiger S, Bultmann U, Schoch B, Gizewski ER, Timmann D, Maschke M, Frings M (2010) Recovery of upper limb function after cerebellar stroke: lesion symptom mapping and arm kinematics. Stroke 41: 2191–2200 Doi 10.1161/STROKEAHA.110.583641

28 Kozberg MG, Yi I, Freeze WM, Auger CA, Scherlek AA, Greenberg SM, van Veluw SJ (2022) Blood-brain barrier leakage and perivascular inflammation in cerebral amyloid angiopathy. Brain Commun 4: fcac245 Doi 10.1093/braincomms/fcac245

29 Kutzelnigg A, Faber-Rod JC, Bauer J, Lucchinetti CF, Sorensen PS, Laursen H, Stadelmann C, Bruck W, Rauschka H, Schmidbauer M et al (2007) Widespread demyelination in the cerebellar cortex in multiple sclerosis. Brain Pathol 17: 38–44 Doi 10.1111/j.1750-3639.2006.00041.x

30 Kyriakou EI, van der Kieft JG, de Heer RC, Spink A, Nguyen HP, Homberg JR, van der Harst JE (2016) Automated quantitative analysis to assess motor function in different rat models of impaired coordination and ataxia. J Neurosci Methods 268: 171–181 Doi 10.1016/j.jneumeth.2015.12.001

31 Lee NJ, Ha SK, Sati P, Absinta M, Luciano NJ, Lefeuvre JA, Schindler MK, Leibovitch EC, Ryu JK, Petersen MA et al (2018) Spatiotemporal distribution of fibrinogen in marmoset and human inflammatory demyelination. Brain 141: 1637–1649 Doi 10.1093/brain/awy082

32 Liang X, Gong M, Wang Z, Wang J, Guo W, Cai A, Yang Z, Liu X, Xu F, Xiong W et al (2024) LncRNA TubAR complexes with TUBB4A and TUBA1A to promote microtubule assembly and maintain myelination. Cell Discov 10: 54 Doi 10.1038/s41421-024-00667-y

33 Magaki S, Tang Z, Tung S, Williams CK, Lo D, Yong WH, Khanlou N, Vinters HV (2018) The effects of cerebral amyloid angiopathy on integrity of the blood-brain barrier. Neurobiol Aging 70: 70–77 Doi 10.1016/j.neurobiolaging.2018.06.004

34 Marzan DE, Brugger-Verdon V, West BL, Liddelow S, Samanta J, Salzer JL (2021) Activated microglia drive demyelination via CSF1R signaling. Glia 69: 1583–1604 Doi 10.1002/glia.23980

35 Matsuda S, Giliberto L, Matsuda Y, Davies P, McGowan E, Pickford F, Ghiso J, Frangione B, D’Adamio L (2005) The familial dementia BRI2 gene binds the Alzheimer gene amyloid-beta precursor protein and inhibits amyloid-beta production. J Biol Chem 280: 28912–28916 Doi 10.1074/jbc.C500217200

36 Matsuda S, Giliberto L, Matsuda Y, McGowan EM, D’Adamio L (2008) BRI2 inhibits amyloid beta-peptide precursor protein processing by interfering with the docking of secretases to the substrate. J Neurosci 28: 8668–8676 Doi 10.1523/JNEUROSCI.2094-08.2008

37 Matsuda S, Matsuda Y, Snapp EL, D’Adamio L (2011) Maturation of BRI2 generates a specific inhibitor that reduces APP processing at the plasma membrane and in endocytic vesicles. Neurobiol Aging 32: 1400–1408 Doi 10.1016/j.neurobiolaging.2009.08.005

38 Matsuda S, Tamayev R, D’Adamio L (2011) Increased AbetaPP processing in familial Danish dementia patients. J Alzheimers Dis 27: 385–391 Doi 10.3233/JAD-2011-110785

39 McAleese KE, Graham S, Dey M, Walker L, Erskine D, Johnson M, Johnston E, Thomas AJ, McKeith IG, DeCarli C et al (2019) Extravascular fibrinogen in the white matter of Alzheimer’s disease and normal aged brains: implications for fibrinogen as a biomarker for Alzheimer’s disease. Brain Pathol 29: 414–424 Doi 10.1111/bpa.12685

40 Merlini M, Rafalski VA, Rios Coronado PE, Gill TM, Ellisman M, Muthukumar G, Subramanian KS, Ryu JK, Syme CA, Davalos D et al (2019) Fibrinogen Induces Microglia-Mediated Spine Elimination and Cognitive Impairment in an Alzheimer’s Disease Model. Neuron 101: 1099–1108 e1096 Doi 10.1016/j.neuron.2019.01.014

41 Meszaros L, Riemenschneider MJ, Gassner H, Marxreiter F, von Horsten S, Hoffmann A, Winkler J (2021) Human alpha-synuclein overexpressing MBP29 mice mimic functional and structural hallmarks of the cerebellar subtype of multiple system atrophy. Acta Neuropathol Commun 9: 68 Doi 10.1186/s40478-021-01166-x

42 Neureither F, Ziegler K, Pitzer C, Frings S, Mohrlen F (2017) Impaired Motor Coordination and Learning in Mice Lacking Anoctamin 2 Calcium-Gated Chloride Channels. Cerebellum 16: 929–937 Doi 10.1007/s12311-017-0867-4

43 Perez H, Abdallah MF, Chavira JI, Norris AS, Egeland MT, Vo KL, Buechsenschuetz CL, Sanghez V, Kim JL, Pind M et al (2021) A novel, ataxic mouse model of ataxia telangiectasia caused by a clinically relevant nonsense mutation. Elife 10: Doi 10.7554/eLife.64695

44 Petersen MA, Ryu JK, Chang KJ, Etxeberria A, Bardehle S, Mendiola AS, Kamau-Devers W, Fancy SPJ, Thor A, Bushong EA et al (2017) Fibrinogen Activates BMP Signaling in Oligodendrocyte Progenitor Cells and Inhibits Remyelination after Vascular Damage. Neuron 96: 1003–1012 e1007 Doi 10.1016/j.neuron.2017.10.008

45 Pinter D, Ritchie SJ, Doubal F, Gattringer T, Morris Z, Bastin ME, Del CVHM, Royle NA, Corley J, Munoz Maniega S et al (2017) Impact of small vessel disease in the brain on gait and balance. Sci Rep 7: 41637 Doi 10.1038/srep41637

46 Radmard S, Zesiewicz TA, Kuo SH (2023) Evaluation of Cerebellar Ataxic Patients. Neurol Clin 41: 21–44 Doi 10.1016/j.ncl.2022.05.002

47 Raghavan S, Przybelski SA, Lesnick TG, Fought AJ, Reid RI, Gebre RK, Windham BG, Algeciras-Schimnich A, Machulda MM, Vassilaki Met al (2024) Vascular risk, gait, behavioral, and plasma indicators of VCID. Alzheimers Dement 20: 1201–1213 Doi 10.1002/alz.13540

48 Reijmer YD, Fotiadis P, Martinez-Ramirez S, Salat DH, Schultz A, Shoamanesh A, Ayres AM, Vashkevich A, Rosas D, Schwab K et al (2015) Structural network alterations and neurological dysfunction in cerebral amyloid angiopathy. Brain 138: 179–188 Doi 10.1093/brain/awu316

49 Richard E, Carrano A, Hoozemans JJ, van Horssen J, van Haastert ES, Eurelings LS, de Vries HE, Thal DR, Eikelenboom P, van Gool WA et al (2010) Characteristics of dyshoric capillary cerebral amyloid angiopathy. J Neuropathol Exp Neurol 69: 1158–1167 Doi 10.1097/NEN.0b013e3181fab558

50 Ryu JK, Petersen MA, Murray SG, Baeten KM, Meyer-Franke A, Chan JP, Vagena E, Bedard C, Machado MR, Rios Coronado PE et al (2015) Blood coagulation protein fibrinogen promotes autoimmunity and demyelination via chemokine release and antigen presentation. Nat Commun 6: 8164 Doi 10.1038/ncomms9164

51 Sharma B, Gee M, Nelles K, Cox E, Irving E, Saad F, Yuan J, McCreary CR, Ismail Z, Camicioli Ret al (2022) Gait in Cerebral Amyloid Angiopathy. J Am Heart Assoc 11: e025886 Doi 10.1161/JAHA.121.025886

52 Sharma B, Gee M, Nelles K, Cox E, Subotic A, Irving E, Saad F, McCreary CR, Ismail Z, Camicioli Ret al (2024) Associations between white and grey matter damage and gait impairment in cerebral amyloid angiopathy. Gait Posture 113: 553–560 Doi 10.1016/j.gaitpost.2024.08.078

53 Shiotsuki H, Yoshimi K, Shimo Y, Funayama M, Takamatsu Y, Ikeda K, Takahashi R, Kitazawa S, Hattori N (2010) A rotarod test for evaluation of motor skill learning. J Neurosci Methods 189: 180–185 Doi 10.1016/j.jneumeth.2010.03.026

54 Shirzadi Z, Schultz SA, Yau WW, Joseph-Mathurin N, Fitzpatrick CD, Levin R, Kantarci K, Preboske GM, Jack CR, Jr., Farlow MR et al (2023) Etiology of White Matter Hyperintensities in Autosomal Dominant and Sporadic Alzheimer Disease. JAMA Neurol 80: 1353–1363 Doi 10.1001/jamaneurol.2023.3618

55 Shirzadi Z, Yau WW, Schultz SA, Schultz AP, Scott MR, Goubran M, Mojiri-Forooshani P, Joseph-Mathurin N, Kantarci K, Preboske G et al (2022) Progressive White Matter Injury in Preclinical Dutch Cerebral Amyloid Angiopathy. Ann Neurol 92: 358–363 Doi 10.1002/ana.26429

56 Stolze H, Klebe S, Petersen G, Raethjen J, Wenzelburger R, Witt K, Deuschl G (2002) Typical features of cerebellar ataxic gait. J Neurol Neurosurg Psychiatry 73: 310–312 Doi 10.1136/jnnp.73.3.310

57 Tamayev R, D’Adamio L (2012) Inhibition of γ-secretase worsens memory deficits in a genetically congruous mouse model of Danish dementia. Mol Neurodegener 7: 19 Doi 10.1186/1750-1326-7-19

58 Tamayev R, D’Adamio L (2012) Memory deficits of British dementia knock-in mice are prevented by Aβ-precursor protein haploinsufficiency. J Neurosci 32: 5481–5485 Doi 10.1523/JNEUROSCI.5193-11.2012

59 Tamayev R, Matsuda S, Arancio O, D’Adamio L (2012) β-but not γ-secretase proteolysis of APP causes synaptic and memory deficits in a mouse model of dementia. EMBO Mol Med 4: 171–179 Doi 10.1002/emmm.201100195

60 Tamayev R, Matsuda S, Giliberto L, Arancio O, D’Adamio L (2011) APP heterozygosity averts memory deficit in knockin mice expressing the Danish dementia BRI2 mutant. EMBO J 30: 2501–2509 Doi 10.1038/emboj.2011.161

61 Tambaro S, Galán-Acosta L, Leppert A, Presto J, Johansson J (2017) BRICHOS - an anti-amyloid chaperone: evaluation of blood-brain barrier permeability of Bri2 BRICHOS. Amyloid 24: 7–8 Doi 10.1080/13506129.2016.1272451

62 Tambini MD, Norris KA, D’Adamio L (2020) Opposite changes in APP processing and human Abeta levels in rats carrying either a protective or a pathogenic APP mutation. Elife 9: Doi 10.7554/eLife.52612

63 Tambini MD, Yao W, D’Adamio L (2019) Facilitation of glutamate, but not GABA, release in Familial Alzheimer’s APP mutant Knock-in rats with increased beta-cleavage of APP. Aging Cell 18: e13033 Doi 10.1111/acel.13033

64 Thal DR, Ghebremedhin E, Rub U, Yamaguchi H, Del Tredici K, Braak H (2002) Two types of sporadic cerebral amyloid angiopathy. J Neuropathol Exp Neurol 61: 282–293 Doi 10.1093/jnen/61.3.282

65 Timotius IK, Roelofs RF, Richmond-Hacham B, Noldus L, von Horsten S, Bikovski L (2023) CatWalk XT gait parameters: a review of reported parameters in pre-clinical studies of multiple central nervous system and peripheral nervous system disease models. Front Behav Neurosci 17: 1147784 Doi 10.3389/fnbeh.2023.1147784

66 Tomidokoro Y, Lashley T, Rostagno A, Neubert TA, Bojsen-Moller M, Braendgaard H, Plant G, Holton J, Frangione B, Revesz T et al (2005) Familial Danish dementia: co-existence of Danish and Alzheimer amyloid subunits (ADan AND Abeta) in the absence of compact plaques. J Biol Chem 280: 36883–36894 Doi 10.1074/jbc.M504038200

67 van der Holst HM, Tuladhar AM, Zerbi V, van Uden IWM, de Laat KF, van Leijsen EMC, Ghafoorian M, Platel B, Bergkamp MI, van Norden AGW et al (2018) White matter changes and gait decline in cerebral small vessel disease. Neuroimage Clin 17: 731–738 Doi 10.1016/j.nicl.2017.12.007

68 van Horssen J, Brink BP, de Vries HE, van der Valk P, Bo L (2007) The blood-brain barrier in cortical multiple sclerosis lesions. J Neuropathol Exp Neurol 66: 321–328 Doi 10.1097/nen.0b013e318040b2de

69 Vejar S, Pizarro IS, Pulgar-Sepulveda R, Vicencio SC, Polit A, Amador CA, Del Rio R, Varas R, Orellana JA, Ortiz FC (2024) A preclinical mice model of multiple sclerosis based on the toxin-induced double-site demyelination of callosal and cerebellar fibers. Biol Res 57: 48 Doi 10.1186/s40659-024-00529-7

70 Vidal R, Barbeito AG, Miravalle L, Ghetti B (2009) Cerebral amyloid angiopathy and parenchymal amyloid deposition in transgenic mice expressing the Danish mutant form of human BRI2. Brain Pathol 19: 58–68 Doi 10.1111/j.1750-3639.2008.00164.x

71 Vidal R, Frangione B, Rostagno A, Mead S, Révész T, Plant G, Ghiso J (1999) A stop-codon mutation in the BRI gene associated with familial British dementia. Nature 399: 776–781 Doi 10.1038/21637

72 Vidal R, Revesz T, Rostagno A, Kim E, Holton JL, Bek T, Bojsen-Moller M, Braendgaard H, Plant G, Ghiso Jet al (2000) A decamer duplication in the 3’ region of the BRI gene originates an amyloid peptide that is associated with dementia in a Danish kindred. Proc Natl Acad Sci U S A 97: 4920–4925 Doi 10.1073/pnas.080076097

73 Yao W, Yin T, Tambini MD, D’Adamio L (2019) The Familial dementia gene ITM2b/BRI2 facilitates glutamate transmission via both presynaptic and postsynaptic mechanisms. Sci Rep 9: 4862 Doi 10.1038/s41598-019-41340-9

74 Yin T, Yao W, Norris KA, D’Adamio L (2021) A familial Danish dementia rat shows impaired presynaptic and postsynaptic glutamatergic transmission. J Biol Chem 297: 101089 Doi 10.1016/j.jbc.2021.101089

75 Yin T, Yesiltepe M, D’Adamio L (2024) Functional BRI2-TREM2 interactions in microglia: implications for Alzheimer’s and related dementias. EMBO Rep: Doi 10.1038/s44319-024-00077-x

76 Zhu X, Hatfield J, Sullivan JK, Xu F, Van Nostrand WE (2020) Robust neuroinflammation and perivascular pathology in rTg-DI rats, a novel model of microvascular cerebral amyloid angiopathy. J Neuroinflammation 17: 78 Doi 10.1186/s12974-020-01755-y

