## Supplementary figures and images for "Pathological Mechanisms of Motor Dysfunction in Familial Danish Dementia: Insights from a Knock-In Rat Model"

### Supplementary figures

**Supplementary Fig. 1**

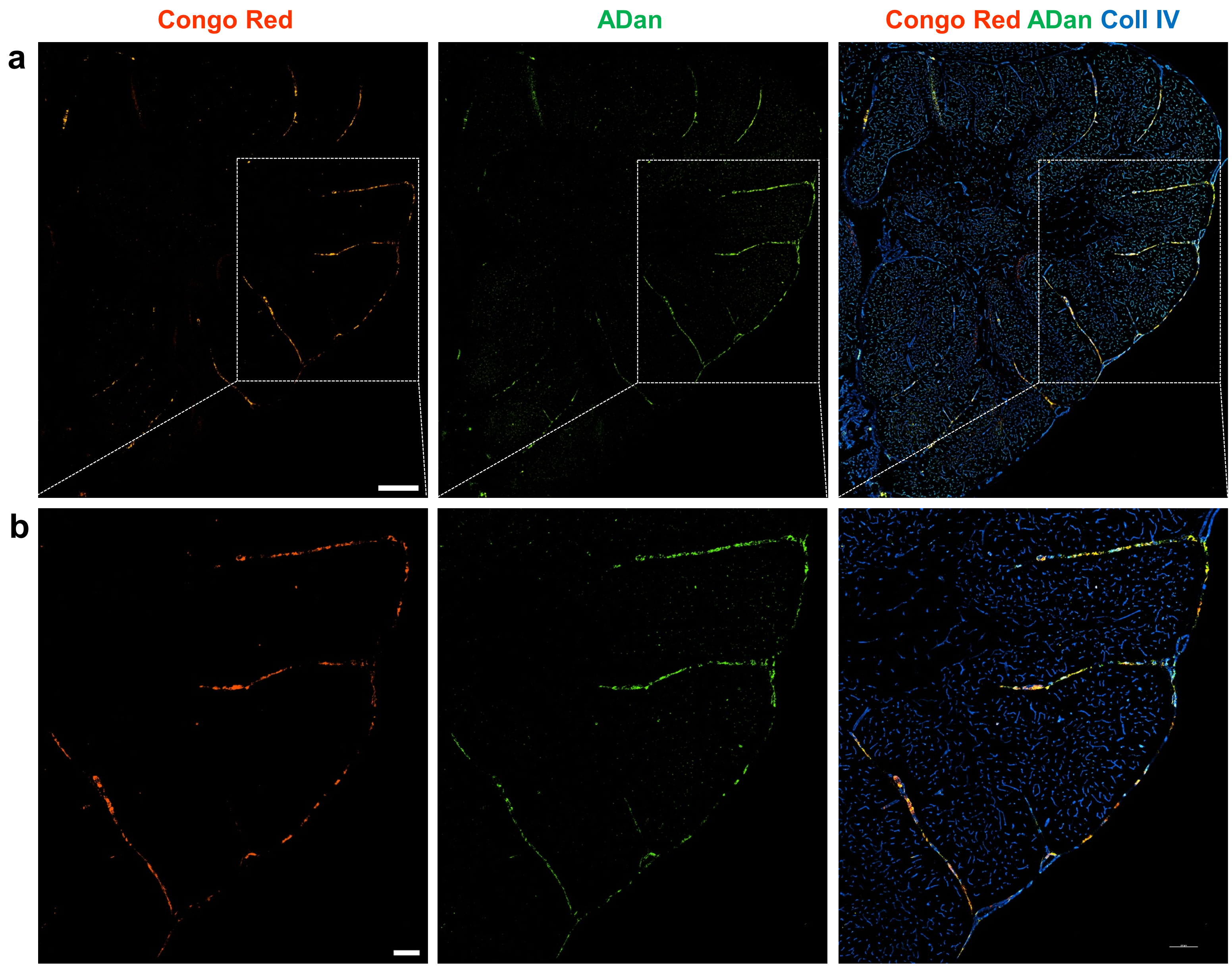

**Supplementary Fig. 2**

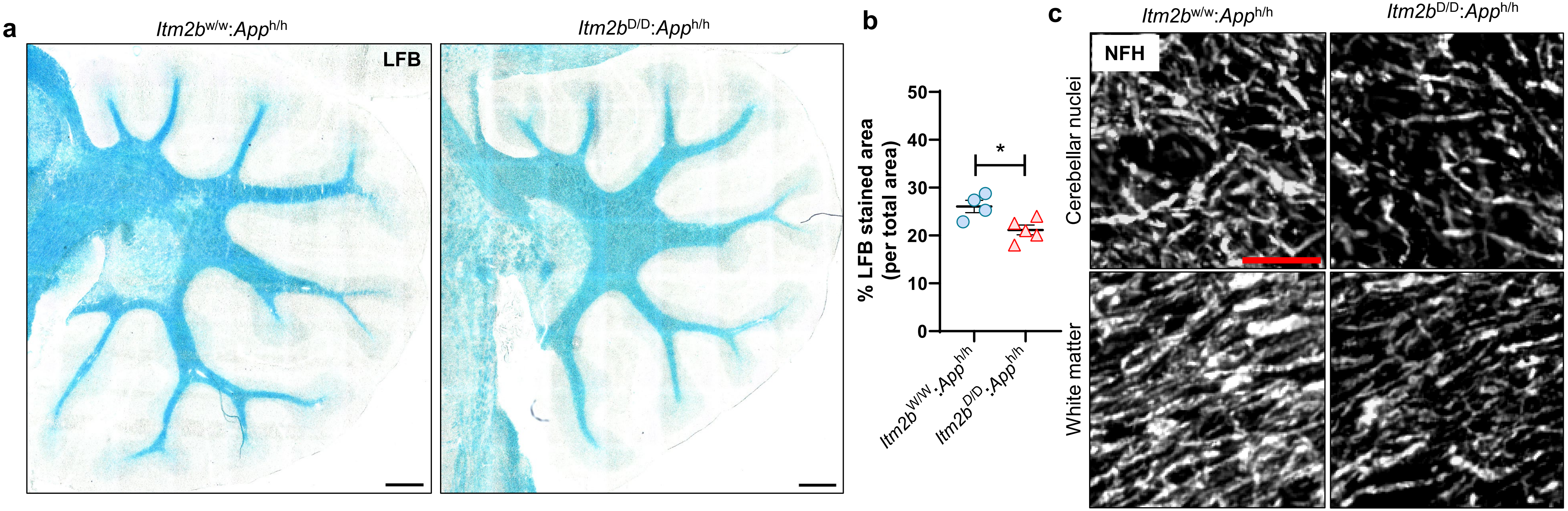

**Supplementary Fig. 3**

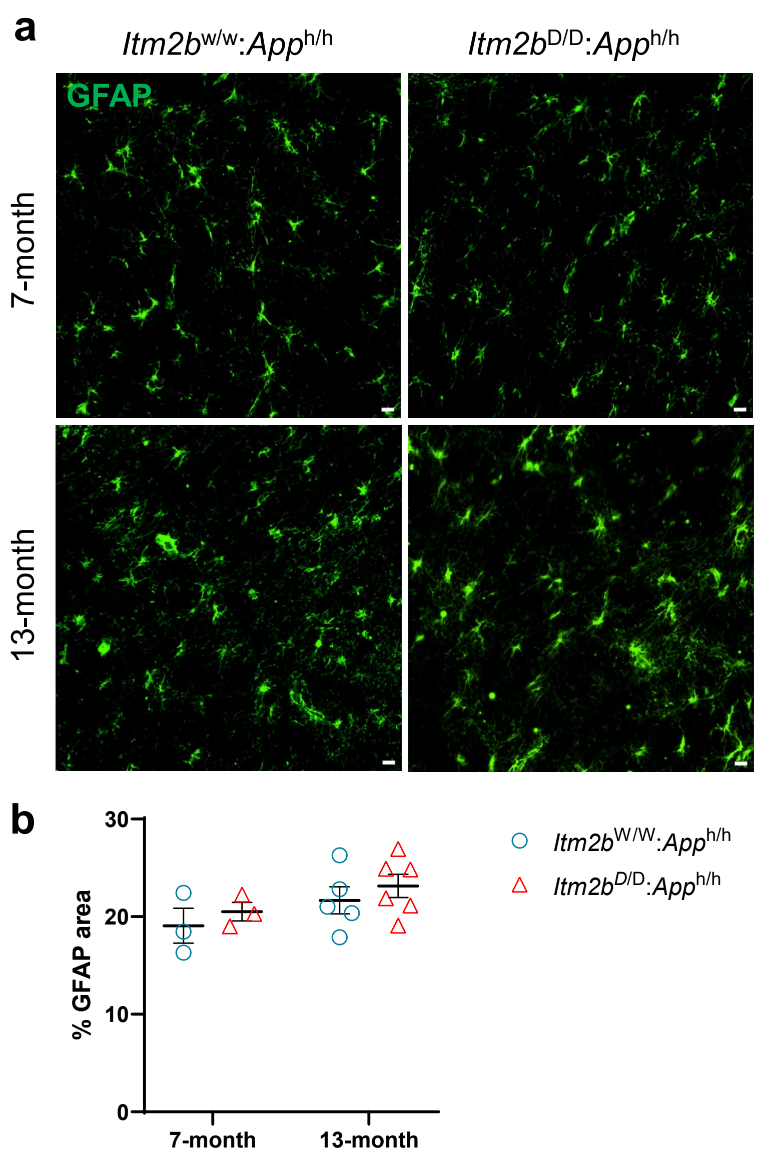
